# Autonomous termination of proliferation in the Drosophila wing disc is TORC1 dependent

**DOI:** 10.1101/2020.12.11.420828

**Authors:** Katrin Strassburger, Marilena Lutz, Sandra Müller, Aurelio A. Teleman

## Abstract

Cells in a developing organ stop proliferating when the organ reaches a correct, final size. The underlying mechanisms are not understood. Although many signaling pathways and cell cycle components are required to sustain cell proliferation, which one of these turns off to terminate proliferation is not known. Here we study proliferation termination using Drosophila wing discs. We extend larval development to provide wing discs a constant growth-sustaining environment, allowing them to terminate proliferation autonomously. We find that the wing pouch, which forms the adult wing blade, terminates proliferation in the absence of brinker or warts, indicating that neither Dpp signaling nor Hippo/Yorkie signaling control final wing size. Instead, termination of proliferation coincides with reduced TORC1 activity and is bypassed by reactivating TORC1. Hence proliferation ceases due to reduced cell growth. Experimental manipulation of Dpp or Yki signaling can bypass proliferation termination in hinge and notum regions, suggesting that the mechanisms regulating proliferation termination may be distinct in different regions of the disc.

**One Sentence Summary:** Using Drosophila, Strassburger et al. investigate the termination of proliferation of an organ when it reaches its final size, and show this occurs due to a drop in TORC1 signaling.

## Main Text

During animal development, organs grow until they reach a suitable final size, at which point the cells in that organ stop proliferating. The correct termination of organ growth is critical for yielding animals of appropriate dimensions and proportions, and for preventing uncontrolled cell proliferation as in cancer (*1*). Although we understand much about the biological mechanisms that control organ patterning, we know comparatively little about how organ size is controlled (*2, 3*).

This fundamental question has been studied mainly in *Drosophila* using the developing wing anlage, the wing disc, which is initially specified as a group of 30 cells that proliferate to yield an organ comprising 50,000 cells at the end of larval development (*2, 4*). At that point, a spike in ecdysone hormone causes animals to stop feeding, become pupae, and undergo metamorphosis, causing growth in the animal to cease (*5–7*). Wing disc cells then perform two final rounds of reductive cell divisions and exit the cell cycle roughly 24 hours after pupa formation (*8–10*).

As wing discs grow during larval development, the cells undergo two separable processes: they grow (increase in biomass) and they proliferate (advance through the cell cycle and divide). Cell cycle progression *per se* does not lead to mass increase (*11, 12*). Hence the cells need to grow. Since diploid cells can only increase in size within a limited range, however, the 1000-fold increase in wing disc size that occurs during larval development requires a combination of cell growth and proliferation. In sum, final wing size is set when wing cells stop growing and proliferating.

Termination of growth and proliferation occurs via a double-safe process involving two redundant mechanisms – one extrinsic to the wing disc and one intrinsic. The extrinsic mechanism is mediated by the hormone ecdysone, which plays a dichotomous role. Intermediate levels of ecdysone during larval development promote wing disc proliferation (*3, 13–15*), whereas high ecdysone levels at the onset of metamorphosis cause wing disc cells to arrest in G2 (*10, 15*). Hence this peak of ecdysone causes termination of growth and proliferation in the wing. Indeed, premature ecdysone secretion causes premature termination of growth, yielding small animals (*16*). The existence of a wing disc-intrinsic size-sensing mechanism has been shown by multiple lines of experimentation (*2*): Firstly, classical experiments in the 1970s where wing discs were transplanted from larvae into adult female abdomens showed that these discs first grow and proliferate, and then stop at roughly the correct size (*4*), indicating the presence of a disc-intrinsic mechanism controlling size in a heterologous environment with constant ecdysone signaling. Secondly, genetic ablation of PTTH or its receptor Torso delays the ecdysone pulse that induces metamorphosis by 5 days, yet adult body size increases by only 0.7-fold and not the 1024-fold, as would be expected from the 12-hour doubling time of wing disc cells (*17, 18*). Hence another mechanism is terminating tissue growth. Finally, experiments manipulating the growth rate of wing compartments revealed that indeed size sensing is an autonomous property of disc compartments (*19*). The signaling pathways involved in this disc-autonomous size sensing, however, are not known.

Several possible mechanisms for disc autonomous size-sensing have been ruled-out. Disc cells do not count cell divisions to determine when to stop proliferating. If large fractions of the disc are killed genetically or with x-rays, the remaining cells compensate by proliferating more than usual, yielding normally sized organs (*20–22*). Indeed, accelerating or decelerating cell cycle progression in the wing disc leads to a normally sized organ composed of either more, smaller cells or fewer, larger cells, respectively (*11, 12*). Likewise, a mechanistic model where cells measure time has been excluded by slowing down disc growth using ‘Minute’ mutations and seeing that discs compensate by extending developmental time, thereby achieving a normal size (*14, 19*).

It is important to differentiate between the signaling pathways that influence the rate of cell growth and proliferation while the disc is in a proliferative state, from the one(s) that cause the termination of cell growth and proliferation when the disc has reached its correct size. A large number of signaling pathways have the capacity to alter the rate of cell growth and proliferation if experimentally manipulated, such as insulin/IGF, Hippo/Yorkie (hpo/yki), Dpp, EGF, JAK/STAT, and Wnt signaling. Which one of these turns off to terminate cell growth and proliferation when an organ has reached its correct size is not known. The pathway responsible for termination should satisfy two criteria. Firstly, the activity of the pathway should change at the timepoint when growth and proliferation cease. If it promotes proliferation, it should become less active. If it blocks proliferation, it should become more active. Secondly, experimentally reverting this change should bypass the termination of proliferation. The pathways which do not satisfy these criteria play a permissive role in allowing cells to grow and proliferate during development, but are not instructive for telling cells when to stop proliferating, and hence are not controlling final organ size.

The Dpp pathway was proposed to regulate wing size. Dpp is secreted from a medial stripe of cells abutting the anterior/posterior compartment boundary. From there Dpp spreads, forming a concentration gradient in the disc (*23, 24*). When experimentally perturbed, Dpp signaling has the capacity to alter cell proliferation and hence organ size (*23, 25–32*). Dpp signaling promotes cell proliferation by inhibiting expression of the transcription factor brinker (brk), which in turn inhibits proliferation by repressing myc and the microRNA bantam (*30, 31, 33–38*). It has been proposed that Dpp signaling needs to increase over time to drive cell proliferation (*39*) however cells that entirely lack Dpp signaling proliferate well if they also lack brk (*37*). It is not known, however, whether Dpp signaling is involved in the termination of wing disc proliferation. To our knowledge, it is not known if Dpp signaling drops when discs terminate proliferating, nor whether wing discs lacking brk bypass this termination. Hence, neither of the two criteria described above are known for Dpp signaling.

The Hippo/Yorkie signaling pathway strongly affects cell proliferation (*40–42*). Cells with elevated Yorkie (yki) activity over-proliferate whereas cells with low yki activity either die or proliferate slowly. Wing discs lacking any upstream negative regulator of yki such as warts (*wts*), salvador, hippo (*hpo*) or mats, massively overgrow (*41, 42*). This occurs, however, when genetic manipulations artificially hyperactivate yki. To our knowledge, it is not known whether yki activity turns off in a wing disc at the end of organ growth. The Hippo pathway senses epithelial tension and cell-cell contacts to induce regenerative proliferation when a tissue is cut or damaged (*40*). Hence the hyperproliferation caused by loss of *wts* or *hpo* may reflect a tissue in an artificial state of constitutive wound healing. Whether this pathway is also involved in termination of proliferation is an open question.

We focus here on the disc-autonomous termination of growth and proliferation of Drosophila wing discs. To do so, we control the disc-extrinsic system by flattening the peak of ecdysone that normally induces metamorphosis, and instead feed ecdysone in a dose-controlled manner to maintain ecdysone signaling within the physiological range of larvae. This allows the larvae to eat and to survive as larvae for many days, enabling us to observe the autonomous termination of wing disc growth and proliferation, and to study which signaling pathways are required for this termination.

We find that a drop in TORC1 signaling, but not yki or dpp signaling, causes the termination of cell proliferation, which is unexpected given that TORC1 mainly promotes cell growth and not cell proliferation (*43, 44*).

## Results

During Drosophila development, ecdysone titers spike above baseline multiple times to promote developmental transitions (Fig. S1A). The spike at the end of larval development induces pupation and metamorphosis. We aimed to flatten this ecdysone peak, which *extrinsically* terminates wing disc proliferation (*10*), thereby extending the larval period and allowing wing discs enough time to terminate growth and proliferation *intrinsically*. To this end, we knocked-down *spookier* (*spok*), an enzyme in the ecdysone biosynthesis pathway (*45, 46*), using a temperature-sensitive inducible system (Tub-Gal4, Tub-Gal80^ts^, hereafter referred to as Tub^ts^>*spok^i^*) at the beginning of the 3^rd^ larval instar (L3) after the animals had completed their last larval molt (Fig. S1A). We monitored ecdysone signaling in whole larvae by measuring mRNA levels of the low-threshold ecdysone target gene *Eip71CD* (also known as *Eip28*/*29*) that is induced by baseline ecdysone levels present in early L3 larvae (*47*). This revealed that ecdysone signaling was reduced within 24 hours of *spok* RNAi induction (Fig. S1B). This inducible system allowed *spok^i^* animals to reach 3^rd^ instar, yet efficiently prevented them from pupating (Fig. S1C). Whereas control animals pupated 72 hours after RNAi induction, and showed increased *Eip71CD* expression, *spok^i^* animals did not (Fig. S1D). Instead, *spok^i^* animals continue feeding and remained active as larvae for an extended period of time. Roughly half of the animals survive as larvae for 2 weeks (Fig. S1E), significantly extending the 3^rd^ instar larval period which usually lasts 2 days.

We next asked what happens to wing disc proliferation in Tub^ts^>*spok^i^* animals, using EdU incorporation as a readout for active progression through S-phase. Although wing discs continued proliferating for the first 12 hours after RNAi induction, they subsequently progressively become EdU negative at 24 and 48 hours after RNAi induction despite still being small (Fig. S1F-F’). This is consistent with our previous findings using tissue explants (*48*), as well findings from other labs (*13, 14*), that some ecdysone signaling is required for wing discs to proliferate. Since *spok* RNAi is reducing ecdysone signaling below the levels of normal L3 larvae (Fig. S1B), this is not sufficient to maintain wing discs in a proliferative state.

To provide wing discs a constant supply of ecdysone at the physiological level of wildtype L3 larvae, we supplemented the food of *spok^i^* animals with the active form of the hormone, 20-hydroxyecdysone (20E). We titrated 3μM to 400μM 20E into the food and found that 25μM 20E was sufficient to restore ecdysone signaling to the level of similarly aged control larvae (Fig. S2A-B). This enabled wing discs to stay proliferative 48 hours after *spok* RNAi induction (Fig. S2C-D, quantified in Fig. S2E-F). In addition to cell cycle progression, 25μM 20E also visibly promoted tissue growth (compare disc size in 0μM vs 25μM Fig. S2C, quantified in more detail below). Similar results were obtained by knocking down *spok* specifically in the prothoracic gland with phm-GAL4 (48h timepoint, Fig. S3). In sum, feeding 25μM 20E to *spok^i^* animals (“Tub^ts^>*spok^i^* +20E”) prevents the ecdysone peak that induces metamorphosis, but, importantly, maintains ecdysone signaling at the physiological level observed in 3^rd^ instar larvae.

Given that Tub^ts^>*spok^i^* +20E wing discs are hosted for many days in a growth-permissive larval environment, analogous to the adult abdomen used in classical transplantation experiments (*4*), we asked whether the wing discs grow and proliferate indefinitely or whether they eventually stop. We assayed growth via incorporation of O-propargyl-puromycin (OPP), which labels nascent polypeptides and is a quantitative readout for translation rates in cells. At 48h after temperature shift in the presence of 25μM 20E, wing discs display incorporation of both EdU and OPP, indicating that they are actively proliferating and growing (Fig. 1A-B’). Interestingly, by 96h they stop proliferating and growing (Fig. 1A-B’). During this time, the wing discs grow circa 10-fold in size and stop at roughly the size of wildtype wL3 discs (Fig. 1C-D). Two possible explanations for the termination of growth and proliferation are: 1) the larval environment is not capable of promoting growth and proliferation beyond 48h or 2) wing discs terminate proliferation because they have reached their final size. To test this, we delayed ecdysone feeding and started it only at 48h after RNAi induction, in order to delay wing disc growth, while larval tissues continue growing. As expected, wing discs of animals where we knockdown *spok* and immediately feed ecdysone are strongly proliferative at 48 hours after RNAi induction and have terminated proliferation 72h after RNAi induction (Fig. 1E-E’, “+20E” samples). Also as expected, wing discs in *spok^i^* animals that are not fed ecdysone are small and non-proliferative at both 48h and 72h after RNAi induction (Fig. 1E-E’, “−20E”). If we take some of these animals and start feeding them ecdysone at 48h, when the discs are still small and EdU negative, they recommence proliferation and growth, and now are strongly EdU positive at 72h, and then terminate proliferation at 96h (Fig. 1E-E’, “48h− → 48h+” samples). This suggests that larvae after the 48h timepoint are still competent to support wing disc proliferation. As additional evidence, presented below, wing discs mutant for *wts* or TSC2 can sustain proliferation beyond these timepoints, and achieve larger final sizes. Furthermore, wing discs that have stopped proliferating do not display cell death (Fig. S4). In sum, wing discs in the Tub^ts^>*spok^i^* +20E system stop growing and proliferating at roughly the right size, despite the lack of an ecdysone pulse that normally stops growth and proliferation extrinsically.

**Figure 1:**
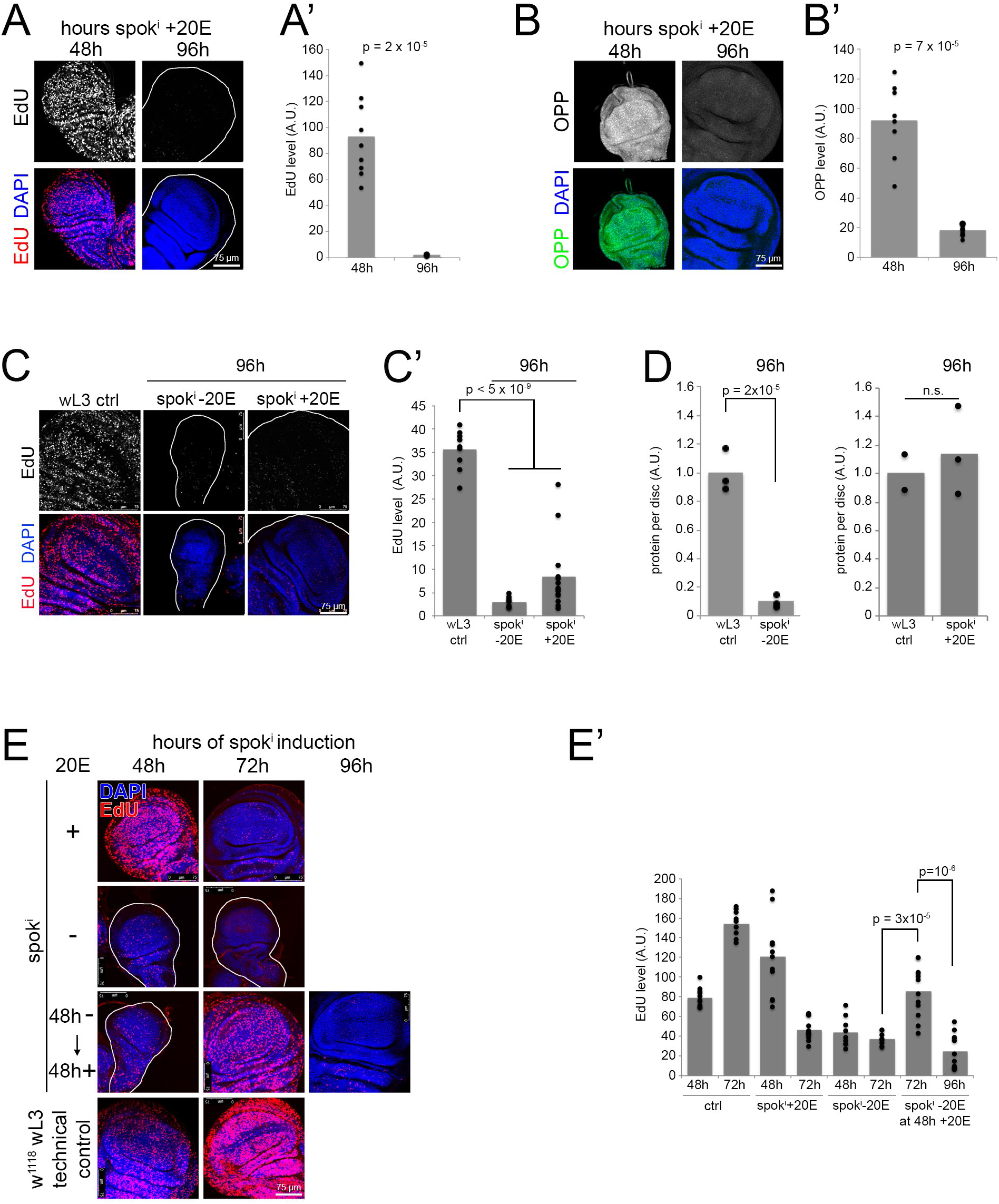
Disc intrinsic termination of proliferation and growth. **(A-B’)** Tub^ts^>*spok^i^* +20E discs first proliferate (A-A’) and grow (B-B’) robustly and then stop. (A-A’) Proliferation, assayed by EdU incorporation, for *spok^i^* discs 48 or 96 hours after knockdown induction and 20E feeding. Representative images in (A), quantified in (A’). n>8 discs/condition x 6 biological replicates. (B-B’) Growth, assayed via O-propargyl-puromycin (OPP) incorporation into nascent polypeptide chains, 48 or 96 hours after knockdown induction and 20E feeding. Representative images in (B), quantified in (B’) n>8 discs/condition x 3 biological replicates. **(C-D)** Tub^ts^>*spok^i^* +20E discs grow 10-fold in size and then terminate proliferation at a size similar to those of control wing discs at the end of 3^rd^ instar larval development. Proliferation is assayed by EdU incorporation, representative images in (C), quantified in (C’) and size is assayed as total protein per disc by BCA protein measurement (D). **(E-E’)** Discs proliferate beyond the 48h timepoint if 20E feeding is delayed. EdU incorporation at indicated times after *spok^i^* induction for larvae not fed ecdysone (−20E), fed ecdysone as of knockdown induction (+20E) or fed ecdysone starting 48h after knockdown induction (48h− → 48h +). The w^1118^ technical control is not grown under the same conditions as the other samples, and only serves to confirm that the EdU staining worked. Representative images in (E), quantified in (E’). n>8 discs/condition x 2 biological replicates.

We tested whether this behavior depends on the concentration of ecdysone being fed to the animals. To this end, we first starved Tub^ts^>*spok^i^* animals of ecdysone for 48h, causing wing discs to stop proliferating (Fig. S1F), and then added 20E to the food. This assures that the signaling and proliferation we subsequently observe are due to the exogenously supplied ecdysone. We titrated 20E in the food from 6μM to 50μM, the maximum possible without inducing pupation. This led to a dose-dependent induction of 20E signaling in the wing disc, assayed by Q-RT-PCR of Eip71CD and Eip74B, which are induced by ecdysone signaling, and ftz-f1 which is inhibited by ecdysone (Fig. S5A). Induction of ecdysone signaling was stable across time, seen by comparing 24h versus 48h after feeding (Fig. S5A). Increasing 20E levels led to a dose-dependent increase in EdU incorporation in the wing disc at 24h (Fig. S5B). In all 20E concentration conditions, however, wing discs terminated proliferation by 48h (Fig. S5B). Hence all the concentrations of 20E we tested were sufficient to support wing disc proliferation, but none was capable of maintaining wing disc proliferation indefinitely. Notably, 20E signaling did not drop in wing discs that had terminated proliferation at 48h compared to 24h (Fig. S5A). Hence wing disc cells do not stop proliferating due to a change in ecdysone signaling. This is conceptually different from situations where ecdysone signaling is experimentally reduced, thereby causing a proliferation block (*13*). Since wing discs stop proliferating without a decrease in ecdysone signaling, some other signal must be causing the proliferation stop.

We next asked what signaling pathways are responsible for terminating proliferation of wing discs. We first assayed whether Dpp signaling turns off when proliferation stops. We stained wing discs for phospho-Mad, which reads-out Dpp signaling activity (*49*), and for brk, which is expressed where Dpp signaling is low, and acts to repress proliferation. This revealed that Dpp signaling is very similar in wing discs 48h after *spok* RNAi induction, when they are small and proliferative, and 96h after *spok* RNAi induction, when growth and proliferation have terminated (Fig. 2A-A’). In both cases, Dpp signaling is high medially, as seen by the high pMad levels, and low laterally where brk is expressed. Hence Dpp signaling does not appear to turn off at the time when proliferation stops. We next tested functionally whether the Dpp/Brk pathway plays a role in termination of proliferation. To this end, we knocked down *brk* by RNAi in the wing disc, since brk mediates the effects of Dpp signaling on cell proliferation (*30, 37*). Wing discs lacking *brk* still terminate proliferation in most regions of the disc, including all of the wing pouch which will form the adult wing blade (Figs. 2B, S6A). Hence brk is not required for termination of proliferation in most regions of the wing disc. In the absence of *brk*, however, wing discs develop lateral bulges, and some wing discs, but not all, show some cell proliferation in these regions at 96h (Figs. 2B, S6A). Similar to the Dpp pathway, the wingless pathway also promotes wing disc growth and proliferation (*32*). The wing pouch of wing discs expressing arm^S10^, which constitutively activates wingless signaling, still terminates proliferating at 96h (Fig. S6B). In sum, although Dpp and wingless signaling can modulate proliferation rates in the wing, neither determines the termination of proliferation.

**Figure 2:**
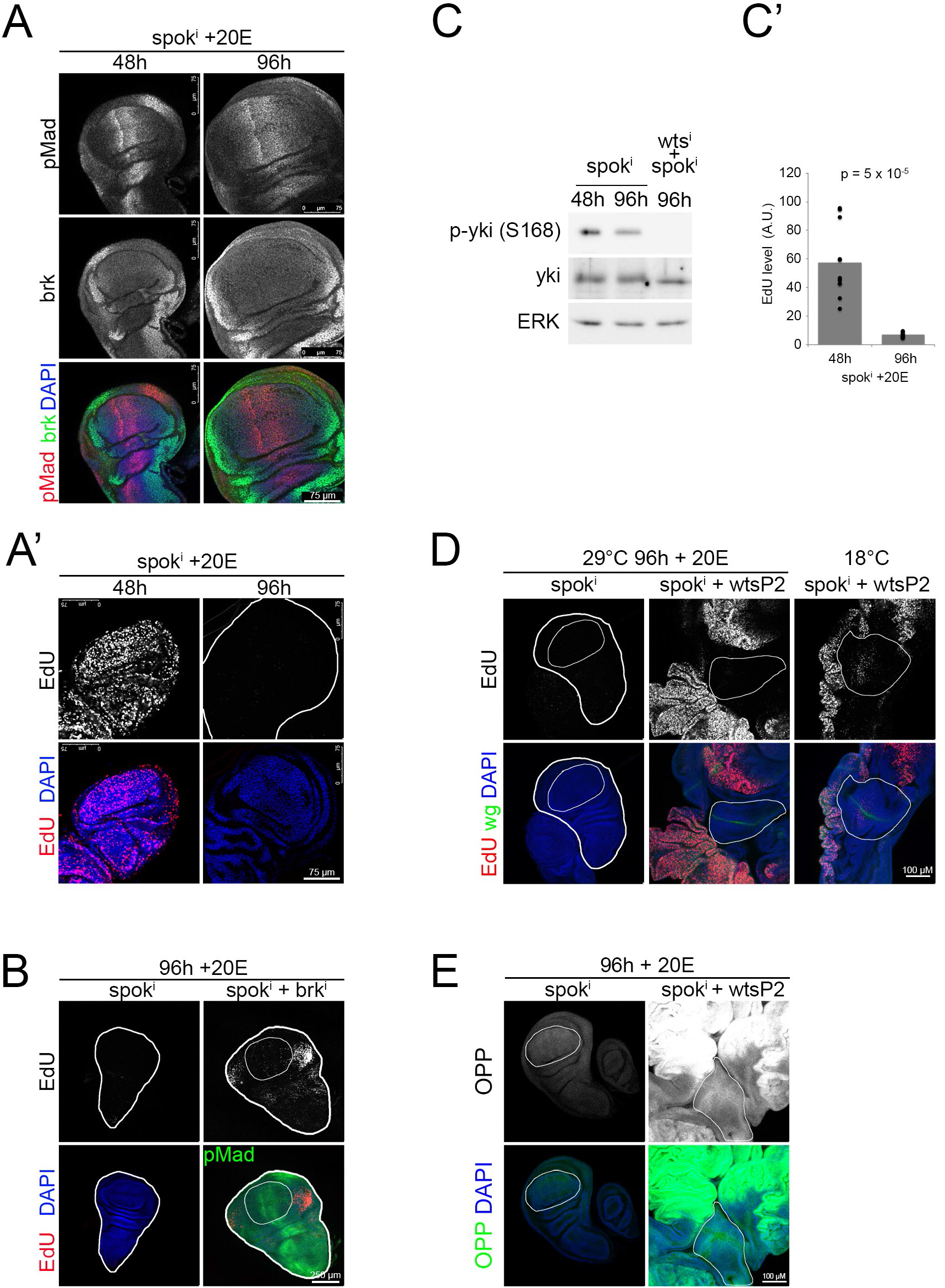
Proliferation in the wing pouch terminates successfully in the absence of *warts* or *brinker*. **(A-A’)** Dpp signaling activity, assayed by immunostaining Tub^ts^>*spok^i^* +20E discs with pMad and brk antibodies (A), is similar in proliferating discs and discs that have terminated proliferation (A’), 48 and 96 hours after knockdown induction, respectively. Proliferation was assayed by EdU incorporation. n=10 discs/condition x 3 biological replicates. **(B)** The wing pouch terminates proliferation also upon knockdown of *brk*. 96h after knockdown induction, Tub^ts^>*spok^i^* +20E discs have terminated proliferation, assayed by EdU incorporation. Most regions of discs with concomitant knockdown of *brk* (*spok^i^* + *brk^i^*) also stop proliferating with only the very lateral regions bypassing the stop. Thick white line outlines the disc, thin white line outlines the pouch. n=10 discs/condition x 3 biological replicates. **(C-C’)** Global yki activity does not drop in Tub^ts^>*spok^i^* +20E discs when they terminate proliferation. (C) Phosphorylation of yki measured by immunoblotting lysates of wing discs 48h or 96h after knockdown induction from Tub^ts^>*spok^i^* +20E or Tub^ts^>*spok^i^* + *wts*^i^ +20E animals. n=40 discs/condition x 3 biological replicates. (C’) Proliferation was measured as a control via EdU incorporation. n=10 discs/condition x 3 biological replicates. **(D)** *Wts* loss of function in Tub^ts^>*spok^i^* +20E discs (*spok^i^* + *wts^P2^*) leads to overproliferation in proximal regions, assayed by EdU incorporation, whereas the pouch (white outline), identified by a *wg* expression stripe, terminates proliferation 96h after knockdown induction and 20E feeding (29°C 96h + 20E). Without spok^i^ knockdown (18°C *spok^i^* + *wts^P2^*) *wts^P2^* discs also stop proliferating in the pouch (white outline) with overproliferating proximal regions. n=10 discs/condition x 3 biological replicates. **(E)** The wing pouch (white outline) of *spok^i^* + *wts^P2^* discs has reduced growth compared to proximal regions, assayed by OPP incorporation, 96h after knockdown induction and 20E feeding. n=6 discs.

We next addressed the role of Hippo/Yorkie (Hpo/Yki) signaling in growth termination. We first tested if yki activity drops when wing discs stop proliferating by quantifying yki phosphorylation with immunoblots of wing disc lysates. Since phosphorylation inactivates yki, we would expect phospho-yki levels to increase when proliferation stops. However, phospho-yki levels did not increase in wing discs 96h after *spok* RNAi induction (Fig. 2C) when the wing discs were no longer proliferative (Fig. 2C’). Nonetheless, we tested functionally whether constitutive activation of yki via knockdown of *warts* (*wts*) can bypass the termination of proliferation. Surprisingly, *wts* knockdown, which very efficiently hyperactivates yki as judged by a strong drop in its phosphorylation (Fig. 2C), has different effects in different regions of the wing disc. The wing pouch, which we identified morphologically (white outline, Fig. S6C), and via the characteristic ZNP double-stripe which is visible by DAPI staining (arrow, Fig. S6C), and by a stripe of *wingless* (*wg*) expression at the D/V boundary, overgrows somewhat but still terminates proliferation in the absence of *wts* (Fig. S6C). This result was unexpected because it indicates that the wing pouch stops proliferating despite active yki. The lateral regions along the D/V boundary – the ones that remain proliferative in the *brk* knockdown discs (Fig. 2B) – also become EdU negative in the absence of *wts* (Fig. S6C). However, both the dorsal and the ventral regions outside the pouch and further away from the D/V boundary remain strongly proliferative in the absence *wts*, and massively overgrow leading to tumor-like masses (example 2, Fig. S6C). OPP incorporation is also lower in the pouch compared to surrounding tissue (Fig. S6D), indicating reduced growth concomitant with the termination of proliferation, and consistent with the size of the wing pouch remaining roughly normal. As a control, we confirmed that wing discs with *wts* knockdown display hallmarks for activated yki: low yki phosphorylation (Fig. 2C) and nuclear yki localization (Fig. S6E). The same results for growth and proliferation were obtained by introducing the wing-disc-specific wts^P2^ mutation, instead of wts-RNAi, into the spok^i^ background (29°C in Fig. 2D, and Fig. 2E). In sum, dpp signaling, wingless signaling, and yorkie signaling do not dictate the termination of growth and proliferation in the wing disc pouch, because signaling through these pathways does not decrease at the timepoint of termination, and because wing pouches terminate growth and proliferation despite experimentally hyperactivated dpp, wingless or yorkie signaling. Hence something else is causing the termination of proliferation.

One regulator of cell growth is TORC1 (*43, 50, 51*). Although TORC1 does not directly regulate cell cycle progression, TORC1 activity is required for cell growth, which in turn is necessary for cell cycle progression (*52*). Immunoblots of lysates from wing discs that had terminated proliferation showed a drop in S6K phosphorylation on Thr398, a direct readout for TORC1 activity (Figs. 3A, S7A). As a consequence, also phosphorylation of S6K on Thr229, the PDK1 site which requires phosphorylation on Thr398 as a priming site, as well as phosphorylation of RpS6 were reduced (Fig. 3A). Phosphorylation of S6K on Thr229 was detected with an antibody that detects the PDK1 phosphorylation sites on both Akt and S6K (Fig. S7B and (*53*)). The drop in TORC1 activity was similar in magnitude to the one obtained by feeding animals rapamycin (Fig. S7C). Consistent with a drop in TORC1 activity, autophagy was mildly induced in ‘stopped’ wing discs (Fig. S7D-D’). Hence, termination of growth and proliferation coincides with a drop in TORC1 activity.

**Figure 3:**
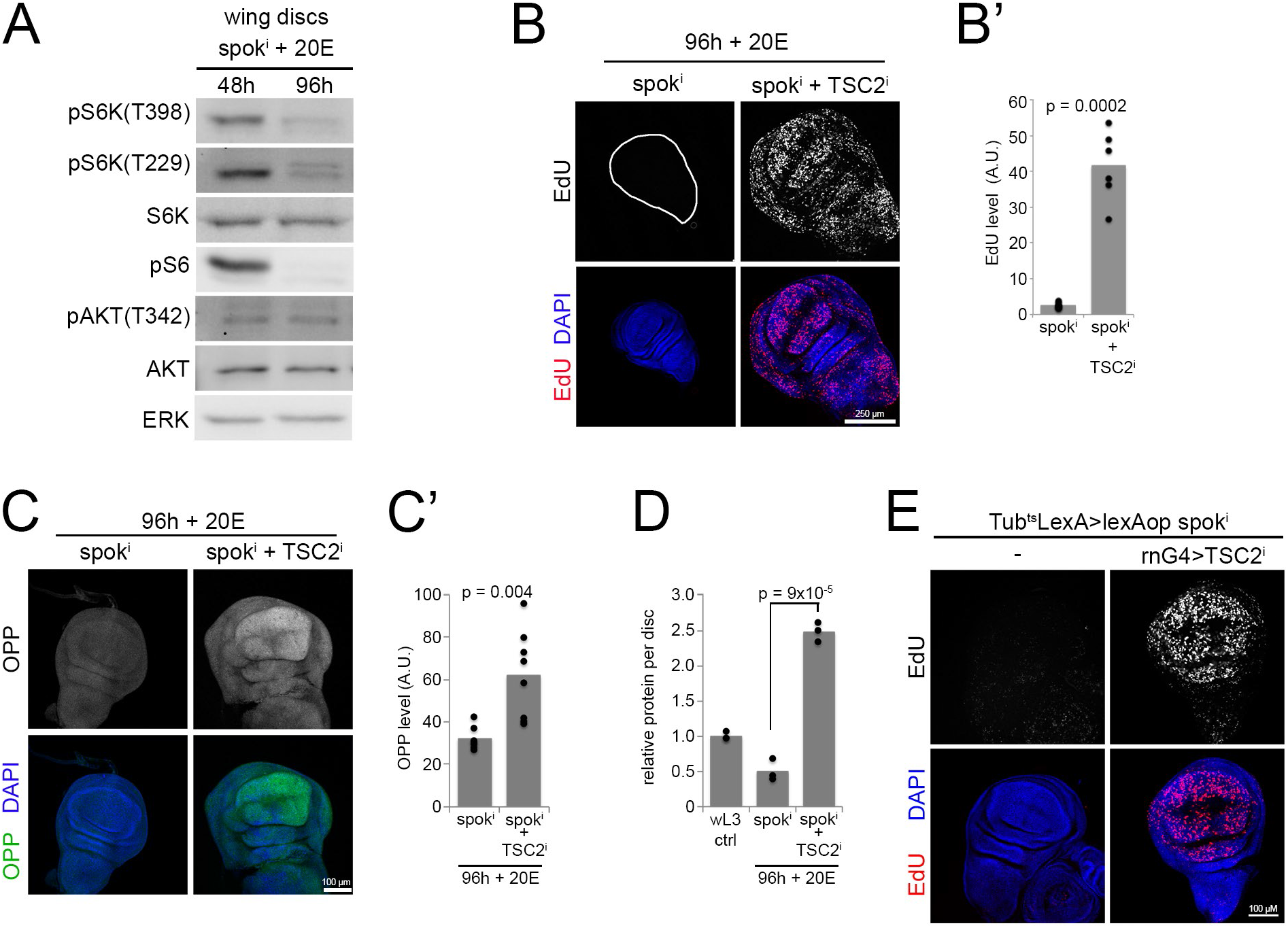
The wing pouch terminates proliferation due to low TORC1 activity. **(A)** TORC1 activity drops when Tub^ts^>*spok^i^* +20E discs terminate proliferation at 96h. Phosphorylation of S6K on Thr398 is a direct readout for TORC1 activity. Phosphorylation on Thr229 by PDK1 requires a priming phosphorylation on Thr398. n=40 discs/condition x 3 biological replicates. **(B-D)** Constitutive activation of TORC1 overcomes the termination of proliferation (B-B’) and growth (C-C’) in Tub^ts^>*spok^i^* +20E discs, causing them to overgrow greatly (D). While Tub^ts^>*spok^i^* +20E discs (*spok^i^* 96h) have terminated proliferation, judged by EdU incorporation, discs with concomitant knockdown of *TSC2* (*spok^i^* + *TSC2^i^* 96h) continue proliferating. Representative images (B and C), quantified in (B’ and C’). n=10 discs/condition x 4 (B-B’) or 3 (C-C’) biological replicates. (D) Disc size at 96h, assayed as total protein per disc by BCA. n=6 discs/condition x 3 biological replicates. **(E)** Wing-pouch specific knockdown of TSC2 bypasses the termination of proliferation in spok^i^ + 20E animals. Spok^i^ is expressed using the lexA/lexAop system combined with Gal80^ts^, and UAS-TSC2^i^ is expressed specifically in the wing pouch with rotund-GAL4 (rnG4). As expected, wing discs stop proliferating in spok^i^ + 20E animals at 96h, and this is bypassed by rnG4>TSC2^i^. n = 13 discs.

To determine whether the decrease in TORC1 activity is upstream or downstream of the termination of cell proliferation, we genetically activated TORC1. If TORC1 inactivation is downstream of a cell cycle block, then re-activating TORC1 should not affect cell proliferation. If, instead, TORC1 inactivation is upstream and causing the cell cycle block, then genetically activating TORC1 should enable wing disc cells to continue proliferating. Ubiquitous activation of TORC1 via knockdown of *TSC2* with Tub^ts^ (Fig. S7A) caused cells in the wing pouch, as well as the rest of the disc, to continue proliferating and growing at 96h when control wing discs stop (Fig. 3B-C’), indicating that the inactivation of TORC1 is upstream of the cell cycle block. *TSC2* knockdown discs become more than twice as large as control discs by 96h (Fig. 3D) and were still proliferating at the latest timepoint we checked, 120h (Fig. S8A-A’). This phenotype was recapitulated with several TSC2 RNAi lines: a VDRC KK line where we recombined away the 2^nd^ insertion at cytological location 40D (Fig. S8B-B’) as well as an independent line from the TRiP collection (Fig. S8C-C’). We confirmed in 3 different ways that this was due to a tissue-autonomous effect of TSC2 knockdown in the wing disc: Knockdown of TSC2 in imaginal discs and the prothoracic gland, using a combination of Ubx-Flp (*54*) and phmG4, also prevented the termination of proliferation (Fig. S9A-A’) whereas knockdown of TSC2 only in the prothoracic gland did not (Fig. S9B-B’). Flip-out clones in the wing disc expressing TSC2 RNAi also bypassed proliferation termination at 96h (Fig. S9C). Finally, we expressed spok^i^ using the lexA/lexAop system and knocked down TSC2 specifically in the wing pouch with rotund-GAL4, and this also led to a bypass of proliferation termination in the wing (Fig. 3E). Since we previously showed that the cell cycle promotes TORC1 via CycD/Cdk4 in the wing disc (*55*), we also tested the other option that TORC1 is downstream of a cell cycle block by overexpressing CycD/Cdk4, however this did not cause cells to continue proliferating (Fig. S9D). In sum, low TORC1 activity is the limiting factor causing the autonomous termination of both growth and proliferation in the wing pouch.

We asked whether our findings hold true in animals where we do not experimentally control ecdysone signaling. We first analyzed wing discs from Tub^ts^>*spok^i^* + wts^P2^ larvae that were not shifted to 29°C, thereby allowing unperturbed endogenous 20E synthesis, and found that also in these discs the pouch terminates proliferating at roughly the correct size while proximal areas continue proliferating, leading to massive overgrowths (18°C, Fig. 2D). Similarly, wing discs from wts^P2^ L3 larvae in an otherwise wildtype background show a range of phenotypes, with some wing pouches completely EdU-negative and some wing pouches containing proliferative cells along the compartment boundaries with EdU-negative quadrants (Fig. 4A). Hence also in this experimental setup the wing pouch can terminate proliferation despite hyperactive yki. In these wts^P2^ mutant discs, TORC1 activity is also low in the pouch (Fig. 4B), indicating that the termination of pouch proliferation coincides with a reduction in TORC1 activity. Knockdown of TSC2 specifically in the pouch of wts^P2^ wing discs significantly increases EdU incorporation in the pouch (Fig. 4C-C’) whereas knockdown of brinker does not (Fig. S9E). In sum, the key findings we present here – that the wing pouch terminates proliferation due to a drop in TORC1 activity but not yki activity – are also observed in a system where ecdysone levels are under physiological regulation.

**Figure 4:**
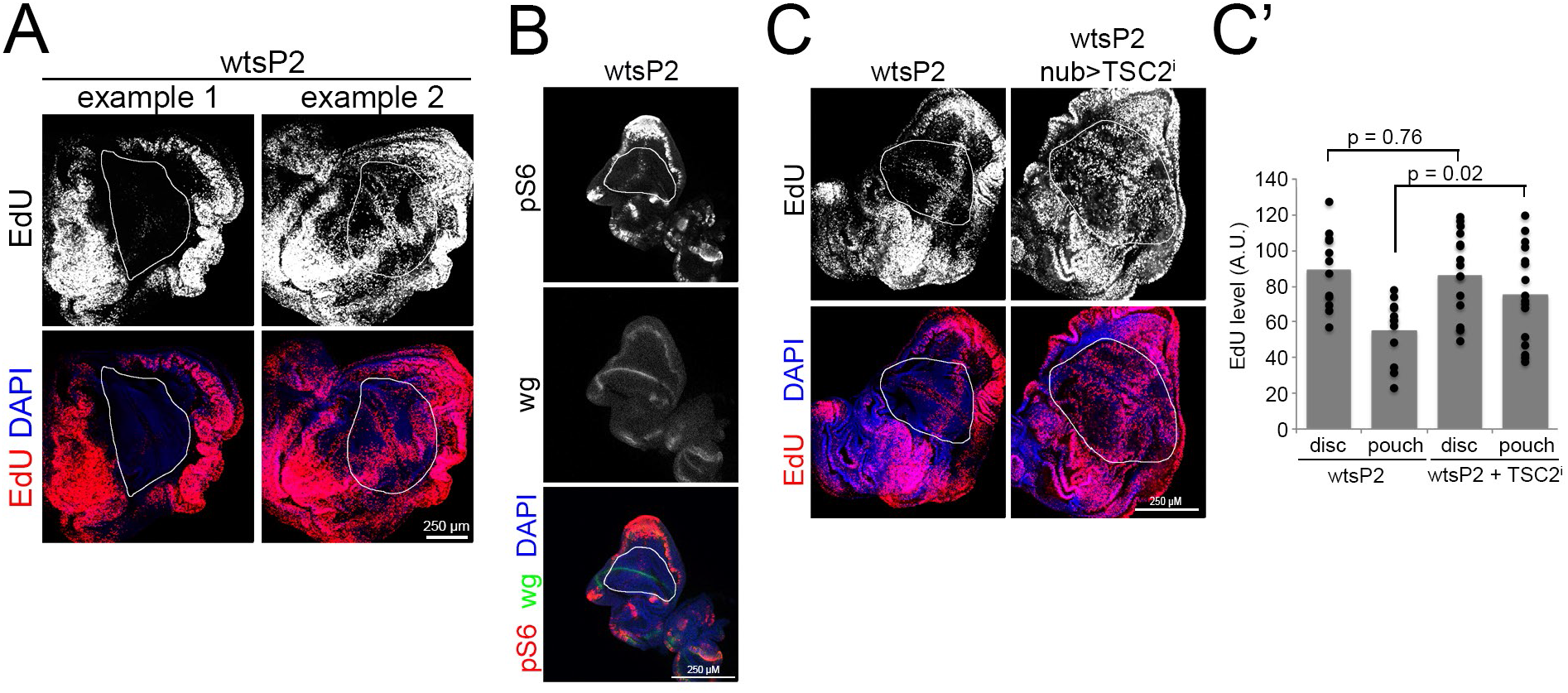
A drop in mTORC1 signaling but not yki signaling terminates pouch proliferation also in animals where ecdysone is under physiological regulation. **(A)** The wing pouch of *wts^P2^* (in WT background) discs can stop proliferating in L3 larvae. A range of phenotypes is observed, from example 1, where the pouch is completely EdU negative, to example 2, where the compartment boundaries are still EdU positive and the quadrants are EdU negative. n=10 discs/condition x 3 biological replicates. **(B)** TORC1 activity drops in the pouch (white outline) of wts^P2^ L3 larvae, measured by immunostaining discs with phospho-S6 antibody. **(C-C’)** Activation of TORC1 in the pouch (white outline) with nubG4 in wts^P2^ L3 wing discs (wts^P2^, nub>TSC2^i^) overcomes the stop of proliferation in the pouch. Representative images (C), quantified in (C’). n = 10 discs x 2 replicates.

Although Akt is one upstream regulator of TORC1, in Drosophila we previously showed that Akt only regulates TORC1 in the ovary, and not other tissues (*56*). In agreement with this, we assayed Akt activity in multiple different ways and found that it does not drop when wing discs terminate growth and proliferation. Phosphorylation of Akt on the PDK1 and TORC2 sites does not decrease (Fig 3A, Fig. S10A), indicating that growth-factor signaling via PI3K is not changing when discs terminate proliferation. If anything, phosphorylation of Akt increases, consistent with a negative feedback loop from TORC1 to PI3K and TORC2. Blotting for all Akt substrates with a pan Akt phospho-substrate antibody on wing disc lysates did not show a drop at 96h (Fig. S10B). Phosphorylation of GSK3b on Ser9, a direct readout for Akt activity, did not drop on a western blot (Fig. S10C) nor by wing disc immunostaining (Fig. S10D-E). Finally, the cytosolic localization of FOXO, which requires Akt activity, also did not change (Fig. S10F). Another upstream regulator of TORC1 is Erk, but Erk phosphorylation also did not change (Fig. S10A). Nutrients signal to TORC1 via the Rag GTPases. However, expression of constitutively active RagC did not bypass the termination of proliferation (Fig. S10G). Hence further work will be required to understand the upstream signals leading to TORC1 inhibition when wing discs reach their target size.

## Discussion

We study here the size sensing mechanism of Drosophila wing discs, focusing on the termination of proliferation and growth. By flattening the peak of ecdysone that induces metamorphosis at the end of larval development, we provide wing discs with a larval environment for an extended period of time of up to 2 weeks. This allows wing discs to terminate proliferation autonomously, rather than as a consequence of the metamorphosis initiated by the ecdysone peak. Important to point out is that the wing discs do not stop proliferating because of an extrinsic limitation of space or nutrients, since *wts* and TSC2 knockdown discs overproliferate massively (Figs. 2D, 3B-D). Also important is that the larval environment is competent to sustain proliferation because the wing discs do proliferate and grow robustly for more than 48 hours before terminating. Hence, the data provided here indicate wing discs have an intrinsic size-sensing mechanism.

We find that the size of the wing blade is determined by a drop in TORC1 activity. Termination of proliferation coincides with a drop in TORC1 activity, and is bypassed by reactivating TORC1. Cell growth and cell cycle progression are intertwined, so that cells usually stop growing if they stop proliferating, and the other way around. Interestingly, the finding that a drop in TORC1 activity controls the termination of proliferation implies that cell cycle progression in the pouch terminates because of a drop in cell growth, and not the other way around. This can happen if TORC1 activity, and hence cell growth, becomes too low for cells to pass the G1/S size checkpoint. This drop in TORC1 activity occurs via an upstream signal which we have not yet identified, but will be the focus of future work.

Unexpectedly, we found that the wing pouch, which gives rise to the adult wing blade, terminates proliferation in the absence of *brk* or *wts*. This indicates that neither Dpp signaling nor Hpo/Yki signaling determine the size at which the wing pouch stops proliferating, and hence the size of the adult wing blade. Do Dpp/Brk and Hpo/Yki signaling control tissue size outside the wing pouch? This is less clear. Both brk loss-of-function as well as yki hyperactivation can cause a bypass in proliferation termination outside the pouch. However, brk is present laterally not only when cells in this region stop proliferating, but also when they are actively proliferating in 3^rd^ instar. Previous reports have shown that lateral brk reduces the rate of proliferation in this region thereby equalizing proliferation throughout the wing disc (*31, 57*). Hence, we favor a model whereby Dpp/Brk signaling differentially modulates the rate of proliferation of different regions of the wing disc to give it a particular shape, but it does not determine when proliferation stops. Regarding yki, we do not see a drop in global yki activity in terminated discs compared to proliferating discs (Fig. 2C). Indeed, a more detailed and spatially-resolved analysis of yki activity in wing discs throughout larval 3^rd^ instar found that yki activity in proximal regions, if anything, increases as animals approach the end of larval development (*58*), which indicates yki is not turning off to terminate proliferation here. Thus future work will be necessary to understand better whether Dpp/Brk or Hpo/Yki signaling regulate growth and proliferation outside the pouch. The mechanisms regulating proliferation termination may be distinct in different regions of the disc.

In sum, we show here that growth and proliferation in the wing pouch, which yields the wing blade, terminate autonomously due to a decrease in TORC1 activity, and future work will be needed to decipher the upstream regulatory mechanisms.

## Materials & Methods

### Fly stains and husbandry

*Spok* knockdown was achieved by crossing Tub-GAL4, Tub-GAL80^ts^/TM6B females to y,w; UAS-*spok* RNAi; UAS-*spok* RNAi males (kind gift from Mike O’ Connor). To collect experimental animals, flies were allowed to lay eggs in vials for roughly 8 hours. Tubes were kept at 18°C for 6 days to reach the L2/L3 molt. Non-TM6B, early L3 larvae were manually picked and transferred to vials containing food previously mixed with 20E (25μM final concentration, unless otherwise indicated), and placed at 29°C to induce spok RNAi expression. Every 48 hours larvae were transferred into food supplemented with fresh 20E. AFG clones expressing TSC2 RNAi were generated by heat-shocking animals at 32°C for 45min directly prior to 20E feeding and the 29°C temperature shift. For rapamycin treatment larvae were fed food supplemented with 200μM rapamycin for 4h prior to disc dissection.

Gene knockdowns were achieved using the following KK lines: VDRC ID 101887 (brk), VDRC ID 106174 (wts), VDRC ID 103417 (TSC2), VDRC ID 103703 (Akt), VDRC ID 101538 (GSK3ß), VDRC ID 104369 (S6K) and Bloomington stocks 51789 (brk^i^ Trip), and 31770 (TSC2^i^ Trip). The following fly stocks were kindly provided by the following people: phmG4, Tub-GAL80ts from Kim Rewitz, UAS-arm^S10^ from Michael Boutros, UAS-CycD from Bruce Edgar, RagC S54N from Kun-Liang Guan and Ubx-FLP from Eugenia Piddini. UAS-Cdk4 was from FlyORF (stock 1652).

To knockdown spok using the lexA/lexAop system: Tubulin-LexA flies were a kind gift from Konrad Basler. To clone LexAop-spok^i^, 454nt of the spok coding region was PCR amplified from genomic DNA of UAS-spok-RNAi flies using OKH974 (ggTCTAGAcgtattttatgtgcta) / OKH975 (ccTCTAGAccgagctaaatttct), and this region was cloned twice in inverse orientation as XbaI fragments first into the AvrI site and second into the NheI site of the pWIZ vector, yielding pKH469. The LexAop-hsp70 promotor was PCR amplified from addgene plasmid #26224 using OKH981 (ggatgcatGCCGGGTCCTCAACGACA) / OKH982 (GCCAGTGCCAGTTCCTGAT) and subcloned as a NsiI/BglII fragment into the SbfI/BglII site of pKH469 yielding the LexAop-spok^i^ construct pKH472.

In order to express TSC2^i^ in the tub^*ts*^>spoki + 20E system (Fig. S8A), TSC2^i^ (VDRC stock 103417, which contains two insertions, at 30B and 40D) was recombined with spok^i^. Progeny were screened by genomic PCR for the presence of the TSC2^i^ at 30B and spok^i^, and absence of the TSC2^i^ 40D locus.

For genotyping stocks and recombinant chromosomes, the following oligos were used (forward/reverse):

spok^i^: CAAGCGCAGCTGAACAAG / GGCACACTCGCTGCATAGT
40D: GCCCACTGTCAGCTCTCAAC / TGTAAAACGACGGCCAGT
30B: GCTGGCGAACTGTCAATCAC / TGTAAAACGACGGCCAGT
wtsP2: CTGCCGCCTGTTTTGAC / AGCGGCTGATGTTGAACTG
Gal4: GATTGACTCGGCAGCTCATC / TGGAACCTGACTCGAAGACC
Gal80: TGGGCAATCAAGACACATT / GCAAGGGCCCATTCTACGA
CycD: CAAGCGCAGCTGAACAAG / TCGCCGGTGAGGACATTTG
CDK4: CAAGCGCAGCTGAACAAG / CCGTCGCGCTCCAGAAAC

The genotypes of animals used in all figure panels are indicated in Supplemental Table 1.

### Quantifications

EdU, OPP, or pGSK3b signals were quantified using ImageJ by defining wing disc area via the presence of DAPI/nuclear stain, and then measuring integrated density of the EdU, OPP or pGSK3b signal in this area. Quantification of lysotracker staining was done analogously, except the wing disc was outlined manually. The signal was then normalized to wing disc area. Note that this results in values with arbitrary units, and should not be used to compare signal levels between different experiments since the absolute value is influenced by staining and imaging settings. To measure pouch area, pouches were outlined using ImageJ and the area was quantified using the ‘measure’ tool.

### Generation of stronger UAS-spok RNAi lines

The original UAS-spok^i^ stock kindly provided by Michael O’Connor, contains two UAS-spok^i^ insertions, on II and III, both of which combined are required to efficiently knockdown spok. To generate single UAS-spok RNAi transgenes strong enough to efficiently knocks down *spok*, we first separated the two UAS-spok RNAi transgenes and transposed the insertion on chromosome II to a different locus using delta2-3 transposase. This yielded the stocks UAS-spok RNAi #4/Cyo and UAS-spok RNAi #8/TM6B. Supplemental Table 1 indicates which UAS-spok RNAi insertion is used in which figure panel.

### Immunoblotting and antibodies

Immunoblotting of discs lysates was performed as described previously (*48*). To detect phosphorylation on S6K, wing discs were collected in batches of 10 discs per PCR tube containing Schneider’s medium on ice. Every batch was immediately spun down after disc collection and the disc pellet was kept on dry ice. Once sufficient wing disc batches were collected, they were successively lysed in one and the same lysis buffer by pipetting up and down and transfering the lysate to the next tube. Lysate was directly boiled at 95°C for 5min. Antibodies used are rabbit anti phospho-S6 kinase (T398) (PhosphoSolutions p1705-398) 1:1000, rabbit anti phospho-Akt (Ser505) (Cell signaling 4054) 1:1000, rabbit anti Akt 1:1000 (Cell signaling 9272), rabbit anti phospho-ERK1/2 1:1000 (Cell signaling 4370), rabbit anti ERK (Cell Signaling 9102) 1:2000, rabbit anti phospho-GSK3b (Ser9) (Cell signaling 9336) 1:1000, rabbit anti phospho-Smad1/5 (Ser463/465) (41D10) (Cell signaling 9516) 1:200, mouse anti rat CD2 (Linaris LFA-2) 1:200, rabbit anti phospho-ribosomal protein S6 (*55*) 1:1000, rabbit anti phospho-Akt Thr342 (*53*) 1:1000, rabbit anti phospho (S/T) Akt substrate (Cell signaling 9611) 1:1000, rabbit anti Drosophila cleaved caspase Dcp-1 (Cell signaling 9578) 1:200. Anti phospho-Yorkie and anti total-Yorkie antibodies were kind gifts from Nic Tapon (*59*) and DJ Pan (*60*). Mouse anti wg antibody was obtained from Hybridoma bank (1:200). Guinea pig anti brk (1:200) and guinea pig anti Drosophila S6K (1:2000) were generated by us and described and validated in previous studies (*61*) (*36, 55*).

### Quantitative RT-PCR

Oligos used for quantitative RT-PCR are (forward/reverse):

Eip71CD: GCGTACCACCAGAAGTATAGGT/AGATTCGGCCATGTCAGCAG
Eip74B: ATCGGCGGCCTACAAGAAG/TCGATTGCTTGACAATAGGAATTTC
Ftz-f1: TGCGAGTCCTGCAAGGGATTCTTCA/GCTCGAACAGCCTCTAGCTTCATGC
rp49: GCTAAGCTGTCGCACAAA/TCCGGTGGGCAGCATGTG

### Total Protein quantification of wing discs

Per genotype three biological replicates were analysed. Per biological replicate 6 wing discs were dissected into PCR tubes containing PBS. Discs were briefly spun down and PBS was removed leaving 10μL behind. 10μl of 12M Urea (6M final) was added and discs were lysed by pipetting up and down repeatedly. Disc lysates were split into two technical replicates and to 10μL each 200μl BCA reagent (Thermo Scientific) was added. BCA reaction mix was incubated for 30min at 37°C before measuring OD562.

### EdU and OPP incorporation reactions

Discs were incubated with EdU or OPP in M3 medium, rotating, and processed and quantified as described previously (*48*). For combined EdU and antibody stainings, first EdU (25μM final concentration) was incorporated into discs during 1h rotation in M3 medium. Afterwards, discs were fixed in 4% paraformaldehyde/PBS for 20min at RT. Discs were permeabilized in PBS/0.2% Triton (PBT) and blocked in PBT/0.1% BSA (BBT) for 30min prior to binding of primary antibodies in BBT O/N at 4°C. The next morning discs were washed in BBT for 1h prior to secondary antibody incubation in BBT for 1h at RT. After secondary antibody binding, discs were washed for 15min in PBT before they were transferred to EdU click-IT reaction mix (100μl for 12 discs) for 30min at RT. Afterwards, discs were washed in PBT and nuclei were stained using DAPI before discs were equilibrated in glycerol mounting medium (160ml glycerol, 20ml 10x PBS, 0.8g n-PG, 12ml water).

### Lysotracker staining

Wing discs and fat bodies were incubated in Schneider’s medium containing 50nM lysotracker red DND-99 (Invitrogen L7528) for 5 min with slight agitation protected from light. Discs were mounted in a large volume of Schneider’s medium on glass slides surrounded by other larval tissues such as mouth hooks or brains in order to prevent tissue damage.

### Statistics

All bar graphs show mean values, with error bars representing standard deviation. Statistical significance was determined by two-tailed, unpaired, students t-test. Prior to application of t-tests, normality of the data were tested using the Shapiro-Wilk test and Kolmogorov-Smirnov tests, except for Q-RT-PCR data where normality was assumed.

### Data availability

All data generated or analysed during this study are included in this published article and its supplementary information files.

## Acknowledgements

We thank Michael O’Connor, Kim Rewitz, Michael Boutros, Bruce Edgar, Kun-Liang Guan, and Eugenia Piddini for flies, and Nic Tapon and DJ Pan for anti-Yki and anti-phospho-Yki antibodies. Anti-wg antibody was obtained from the Developmental Studies Hybridoma Bank, created by the NICHD of the NIH and maintained at The University of Iowa, Department of Biology, Iowa City, IA 52242.

## Author contributions

K.S., M.L. and S.M. performed experiments. All authors designed experiments, analyzed data, and wrote the manuscript.

## Competing interests

The authors declare no competing interests.

## Data and materials availability

All data are available in the main text or the supplementary materials.”

---

**Figure S1:**
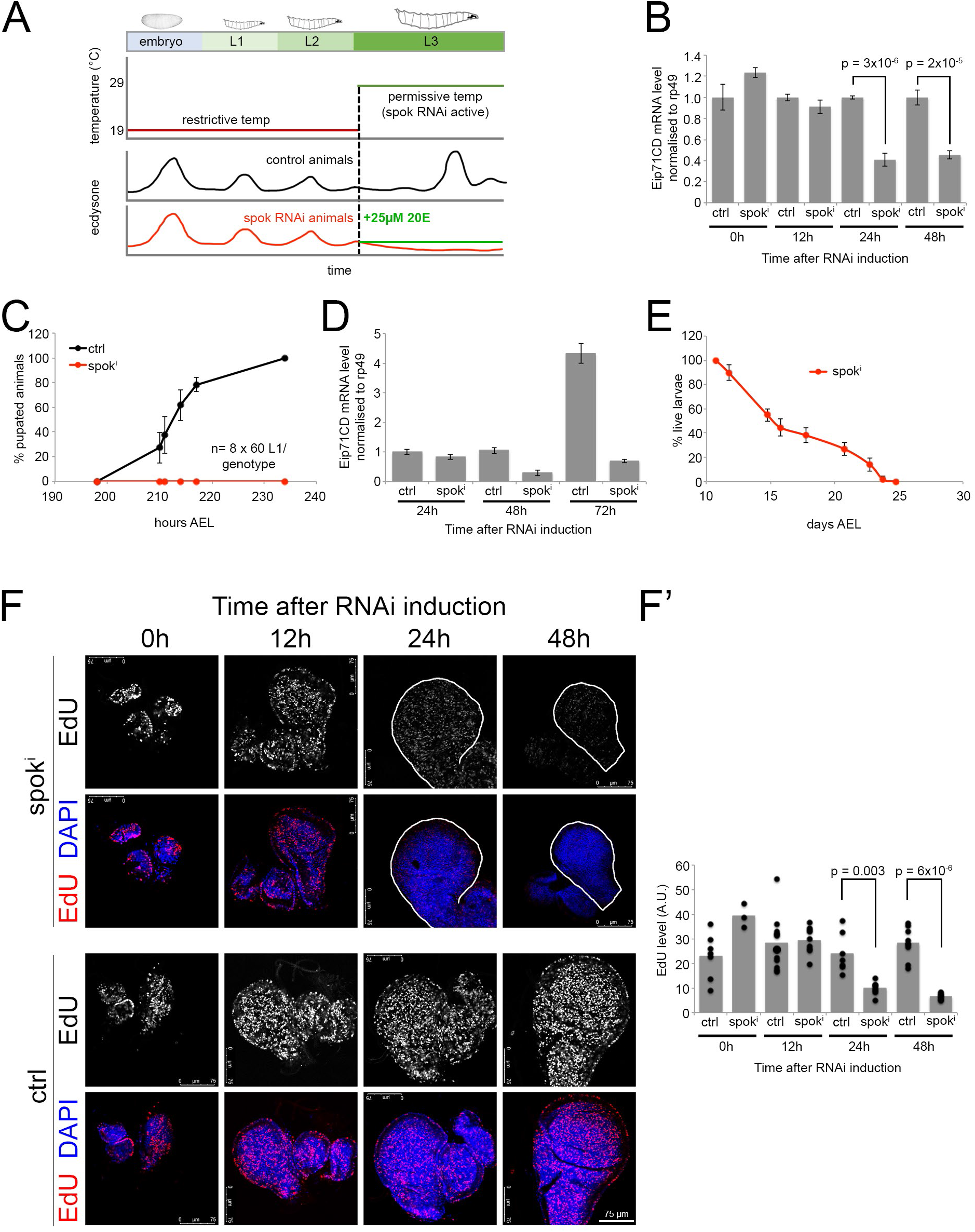
Conditional knockdown of *spok* prolongs larval L3 stage by 2 weeks. **(A)** Schematic representation of the Tub^ts^>*spok^i^* system. Animals are kept at 19°C to allow *spok* expression and normal development until the 3^rd^ instar (L3) stage. After 6 days, after the L2-L3 molt, animals are shifted to 29°C to induce *spok* knockdown, flattening the ecdysone pulse at the end of L3 that induces pupation. Feeding Tub^ts^>*spok^i^* animals 25μM 20E maintains ecdysone signaling in the animal at the physiological level of L3 larvae (green trace). **(B)** Knockdown of *spok* reduces ecdysone signaling. Ecdysone target gene *Eip71CD* mRNA levels, detected by Q-RT-PCR, normalized to *rp49*, from Tub^ts^>+ (ctrl) or Tub^ts^ >*spok* RNAi (*spok^i^*) larvae at 0-48 hours after knockdown induction. n=3 technical replicates x 2 biological replicates x 6 animals/sample. Error bars = std. dev. **(C)** Knockdown of *spok* prevents pupation. The percentage of pupated control (ctrl) or *spok^i^* animals was determined over time. **(D)** Knockdown of *spok* prevents the ecdysone pulse at the end of larval development. *Eip71CD* mRNA levels, detected by Q-RT-PCR, normalized to *rp49*, from control (ctrl) or *spok^i^* larvae at 24-72 hours after knockdown induction. n=3 technical replicates x 2 biological replicates x 6 animals/sample. Error bars=std. dev. **(E)** Tub^ts^>*spok^i^* animals survive as active 3^rd^ instar larvae for several weeks. The percentage of live larvae was counted over time. n=3 technical replicates x 2 biological replicates x 6 animals/sample. Error bars=std.dev. **(F-F’)** Knockdown of *spok* prevents wing disc proliferation. EdU incorporation of control (ctrl) or *spok^i^* wing discs assayed at 0-48h after knockdown induction. Representative images in (F), quantified in (F’) n = 2 biological replicates x >8 discs/condition.

**Figure S2:**
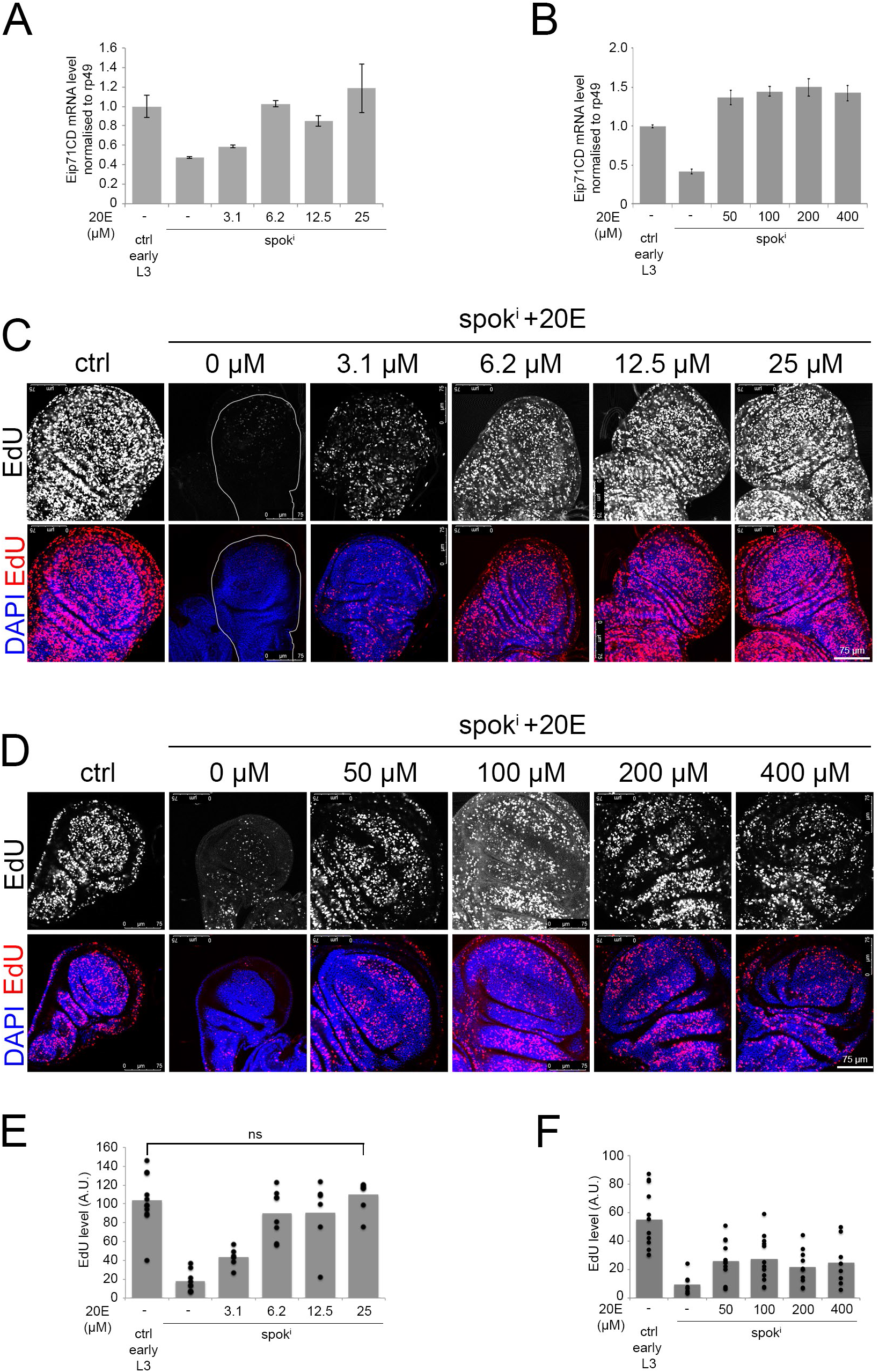
Titration of 20E in the food of Tub>spok^i^ animals identifies a concentration that maintains ecdysone signaling at the physiological level of L3 larvae and supports wing disc proliferation. **(A-B)** 20E supplementation restores the expression of ecdysone target gene *Eip71CD*. *Eip71CD* mRNA levels detected by Q-RT-PCR, normalized to *rp49* for pre-wandering L3 ctrl larvae or larvae 48h after induction of *spok* knockdown with indicated concentrations of 20E supplemented into the food. n=3 technical replicates x 2 biological replicates x 6 animals/sample. Error bars = std. dev. **(C-F)** 20E supplementation restores wing disc proliferation. EdU incorporation 48h after induction of *spok* knockdown and feeding of indicated concentrations of 20E. Representative images in C-D, quantified in E-F. n=3 biological replicates x >8 discs/condition.

**Figure S3:**
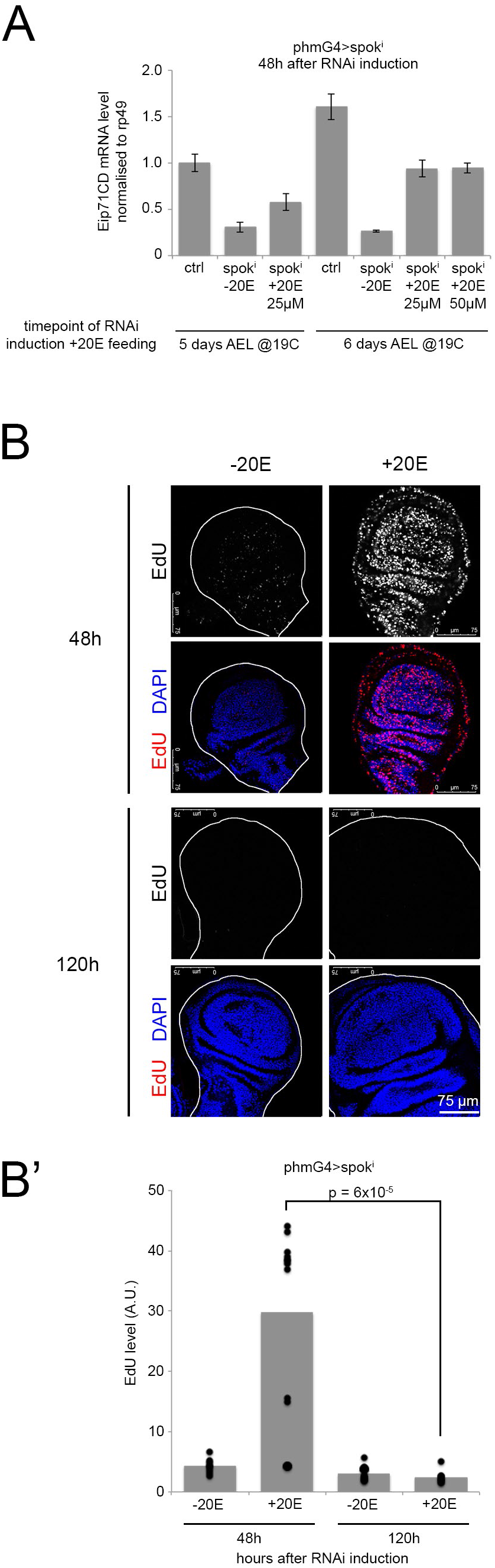
Inducible knockdown of *spok* in the prothoracic gland recapitulates the phenotypes observed upon *spok* knockdown in the entire animal. **(A)** Inducible knockdown of *spok* using phm-GAL4, Tub-GAL80^ts^, UAS-*spok*-RNAi causes reduced ecdysone signaling in larvae 48h after RNAi induction, similar to the Tub^ts^>*spok^i^* system. Animals were grown at 19C, and after 5 or 6 days, as indicated, *spok* RNAi was induced by shifting to 29C. At the same time, larval food was supplemented with the indicated concentrations of 20E. 48 hours after knockdown induction, ecdysone signaling was measured by detecting *Eip71CD* mRNA levels by Q-RT-PCR, normalized to *rp49*. ctrl: Tub>+ 48h after knockdown induction. n=3 technical replicates x 2 biological replicates x 6 animals/sample. Error bars = std. dev. **(B-B’)** Similar to the Tub^ts^>*spok^i^* +20E system, discs in animals with an inducible prothoracic gland-specific knockdown of *spok* (phm^ts^>*spok^i^*) that are fed 20E first proliferate for 48h and then terminate proliferation. By 120 hours after knockdown induction proliferation has stopped as judged by EdU staining. Representative images in (B), quantified in (B’) n=2 biological replicates x >8 discs/condition.

**Figure S4:**
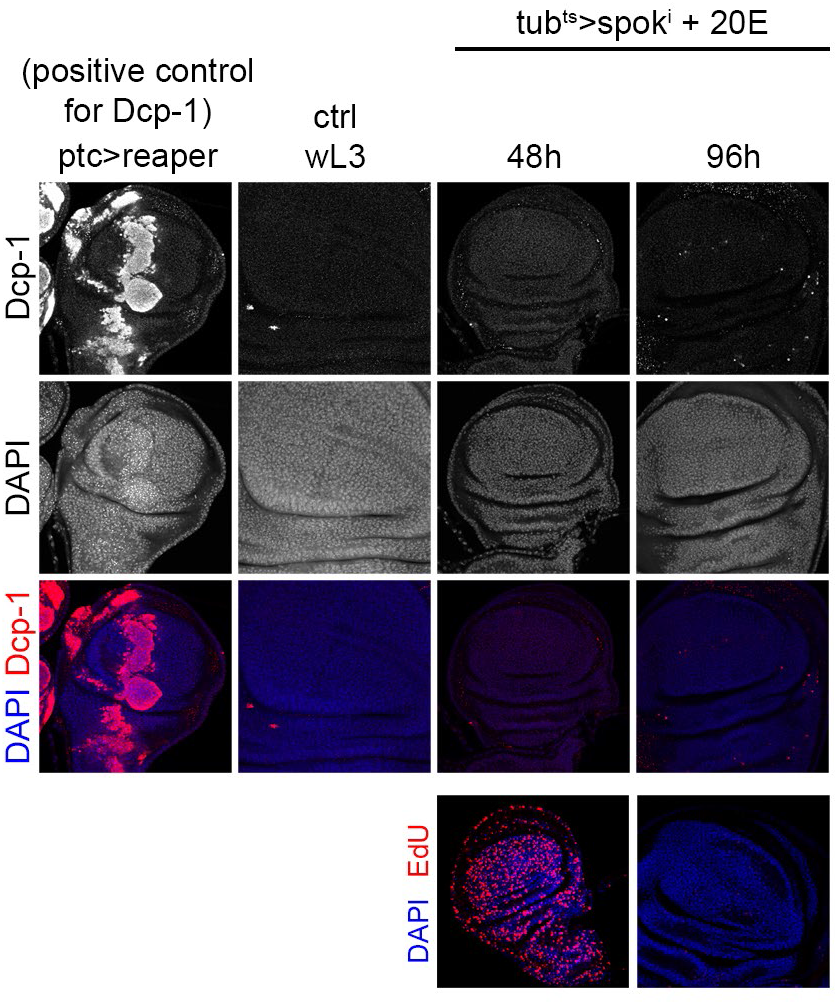
Tub^ts^>*spok^i^* +20E discs are not apoptotic. Cell death in Tub^ts^>*spok^i^* +20E discs was measured by immunostainings with dcp-1 antibody at different timepoints. As a positive control, reaper was expressed for 24h using ptcG4, Tub-GAL80^ts^. Tub^ts^>*spok^i^* +20E discs have a few apoptotic cells, similar to w^1118^ wL3 wing discs which serve as a negative control. n = 8 discs/condition.

**Figure S5:**
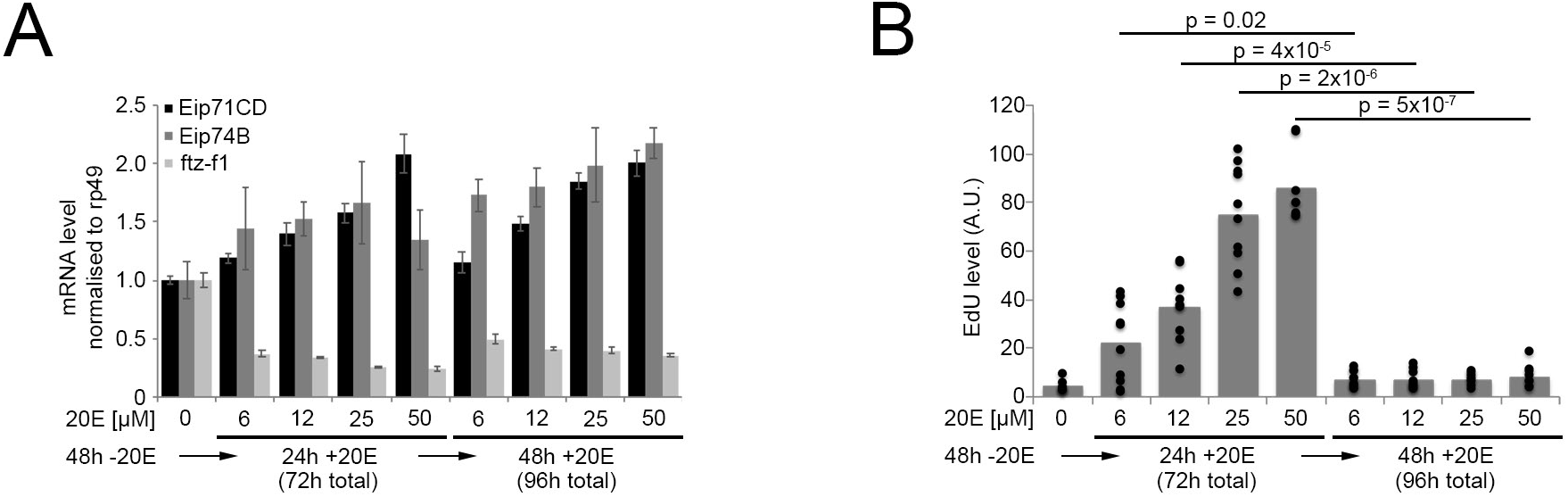
Wing discs stop proliferating at all ecdysone concentrations, and ecdysone signaling does not decrease when wing discs autonomously stop proliferating. Knockdown of *spok^i^* (Tub^ts^>*spok^i^*) was induced 48h prior to feeding larvae with 6-50μM 20E in order to start with a low baseline. (A) Ecdysone target gene expression was measure by Q-RT-PCR 24h and 48h after 20E feeding, whereby Eip71C and Eip74B are induced by 20E and fts-f1 is repressed by 20E. n = 10 discs/condition x 2 biological replicates. Error bars = std. dev. (B) For all ecdysone signaling levels, Tub^ts^>*spok^i^* +20E discs stop proliferating by 48h after 20E supplementation, measured by EdU incorporation. n = 10 discs/condition. Representative of 2 biological replicates.

**Figure S6:**
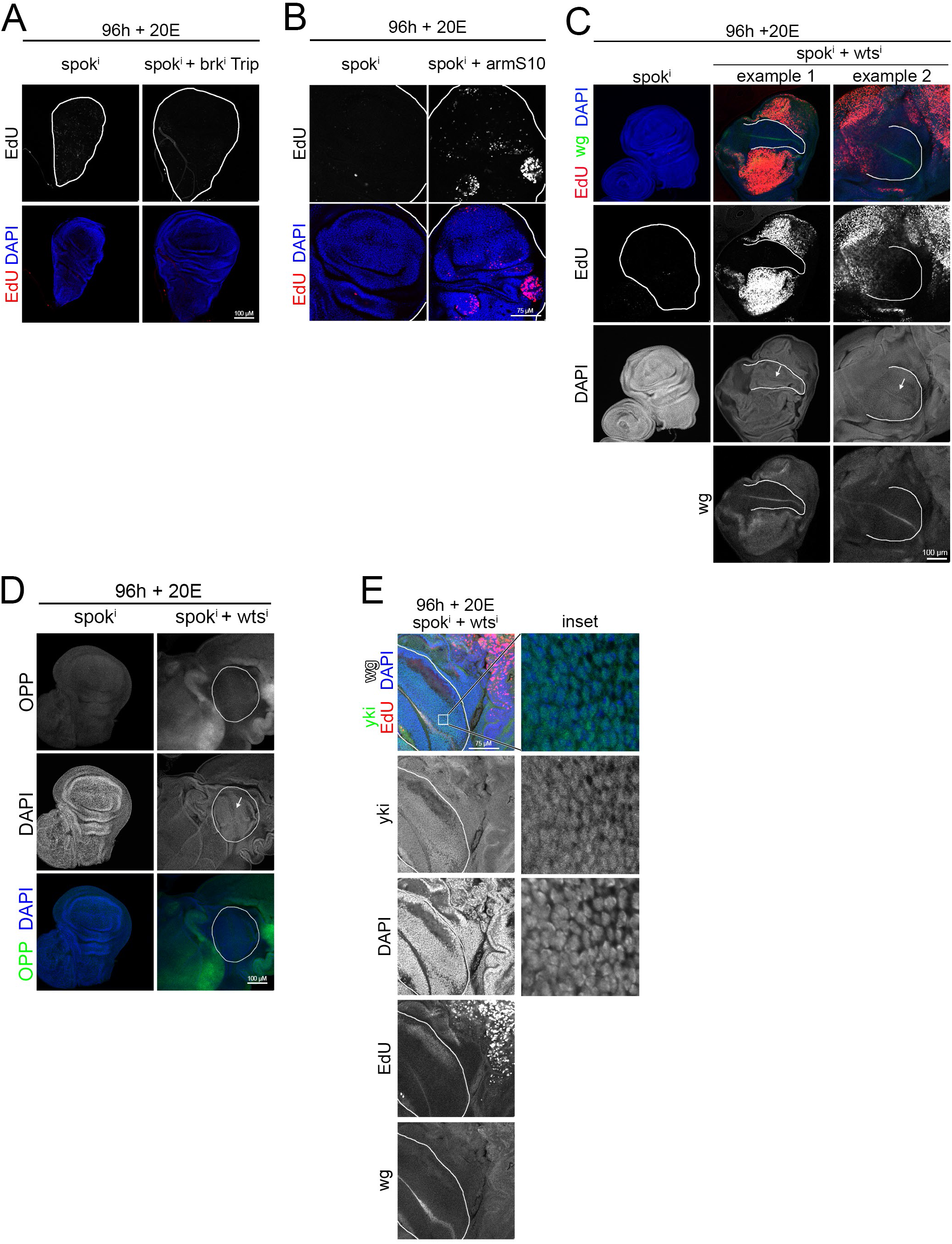
*Dpp*, *hippo* and *wingless* signaling are not part of the mechanism sensing pouch size. **(A)** The wing pouch terminates proliferation despite knockdown of *brk* using a second RNA line (from the TRiP collection) independent of the one shown in main Figure 2b. Both Tub^ts^>*spok^i^* discs and Tub^ts^>*spok^i^*+*brk^i^* discs have terminated proliferation by 96 hours after knockdown induction +20E, assayed by EdU incorporation. n = 16 discs. **(B)** The wing pouch still terminates proliferation upon constitutive activation of wingless signaling by overexpression of armadillo^S10^. The wing pouch of both Tub^ts^>*spok^i^* discs and Tub^ts^>*spok^i^*+*arm^S10^* discs have terminated proliferation by 96 hours after knockdown induction +20E, assayed by EdU incorporation. n = 10 discs. **(C-D)** Knockdown of *wts* in Tub^ts^>*spok^i^* +20E larvae (*spok^i^* + *wts^i^*) leads to overproliferation (C) and overgrowth (D) in proximal regions, assayed by EdU incorporation and OPP incorporation, respectively, whereas the pouch (white outline), identified by the ZNP double-stripe in DAPI staining (arrow) and a *wg* expression stripe, terminates proliferation. The degree of overproliferation in *spok^i^* + *wts^i^* discs varies: two representative examples are shown in (C). **(E)** Knockdown of *wts* causes constitutive activation of yki. Yki localization is mainly nuclear in the wing pouch of *wts*-RNAi wing discs. The wing pouch (white outline) is identified morphologically and marked by the wg expression stripe. n = 10 discs.

**Figure S7:**
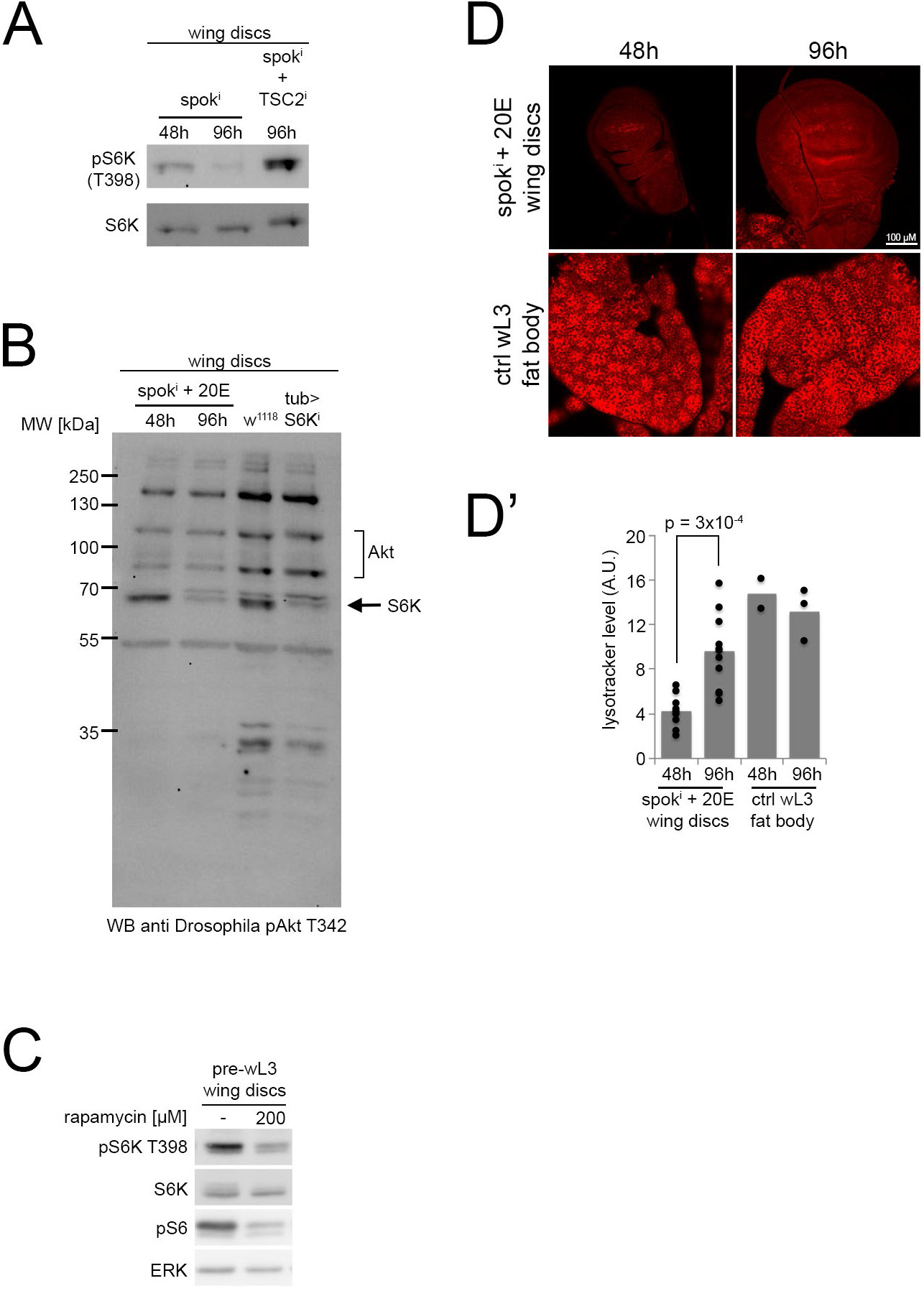
TORC1 activity drops when Tub^ts^>*spok^i^* +20E discs terminate proliferation. **(A)** TORC1 activity drops when Tub^ts^>*spok^i^* +20E discs terminate proliferation, and this is rescued by knockdown of TSC2. Phosphorylation of S6K (Thr398), a direct readout for TORC1 activity, was measured by immunoblotting lysates of wing discs from Tub^ts^>*spok^i^* +20E animals either 48h after knockdown induction, when they are still proliferating, or 96h after knockdown induction when they have terminated proliferation, or from Tub^ts^>*spok^i^* +TSC2^i^ +20E animals at 96h after knockdown induction. n=40 discs/condition x 4 biological replicates. **(B)** Phosphorylation of S6K on Thr229 can be detected using an antibody that detects the very similar PDK1 sites on both Akt and S6K (*53*). 96h after knockdown induction and 20E feeding Tub^ts^>*spok^i^* discs show a drop in pS6K Thr229. The S6K-specific band (arrow) was identified by comparing wing disc lysates of control (w^1118^) and S6K RNAi (Tub>S6K^i^) expressing larvae. **(C)** Feeding larvae rapamycin reduces S6K phosphorylation in wing discs, and shows that the lower band detected by the pS6K antibody does not respond to changes in TORC1 activity within physiological range. Pre-wandering L3 larvae were kept on food containing indicated amounts of rapamycin for 4h before wing disc lysates were subjected to immunoblotting with indicated antibodies. n = 20 discs/condition. **(D-D’)** Consistent with TORC1 activity dropping, autophagy mildly increases in Tub^ts^>*spok^i^* +TSC2^i^ +20E discs 96h after knockdown induction and 20E feeding, measured by lysotracker staining. Representative pictures (D), quantification of lysotracker signal intensity (D’). n = 9 discs. Ctrl wL3 fat bodies were used as positive controls.

**Figure S8:**
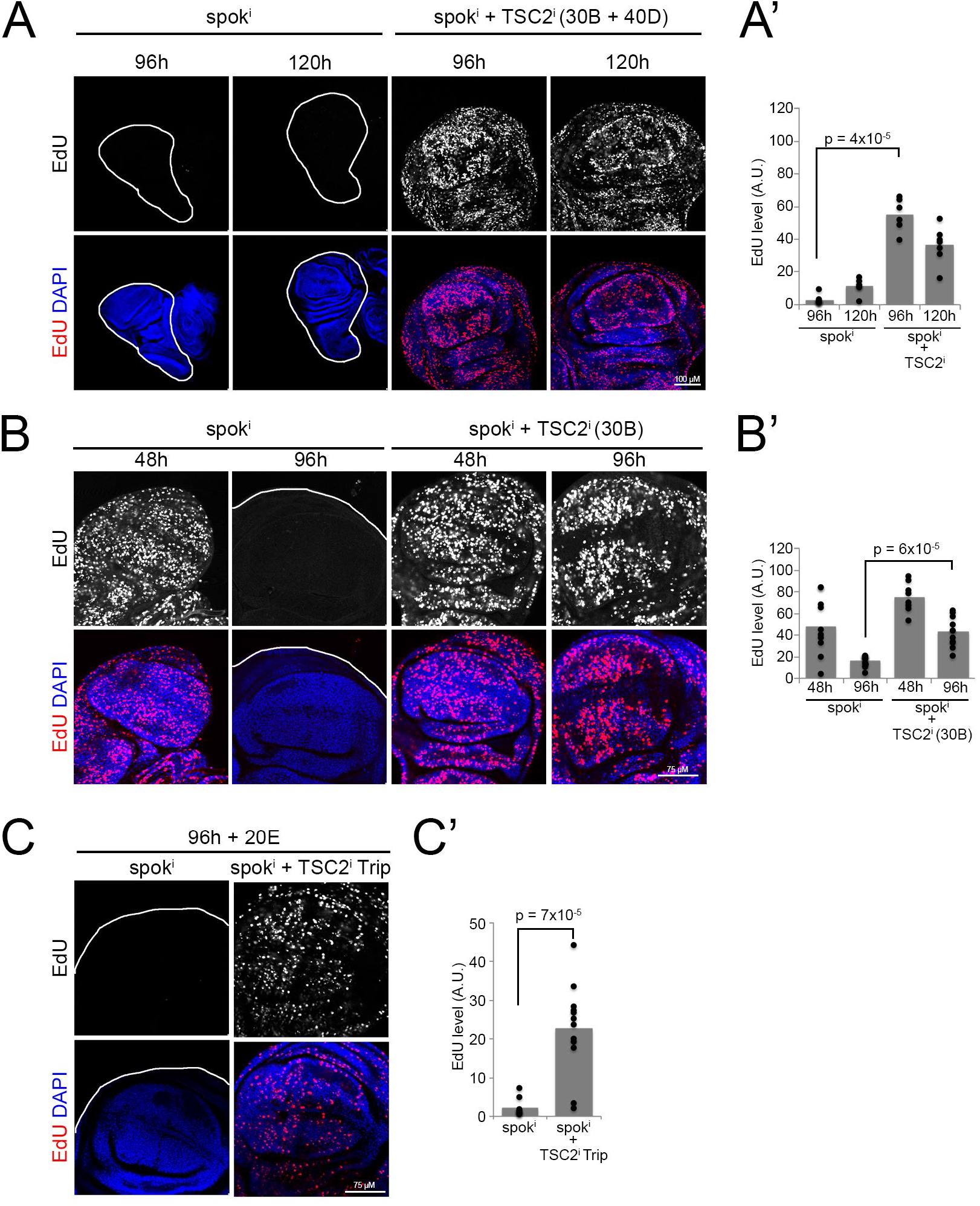
Activation of TORC1 with different RNAi lines against TSC2 prevents proliferation termination in Tub^ts^>*spok^i^* + 20E discs. **(A-A’)** Tub^ts^>*spok^i^* +TSC2^i^ +20E discs stay EdU positive for at least 120h after knockdown induction and 20E feeding. The RNAi lines used here is the original line from the VDRC KK collection (number 103417) which contains two TSC2-RNAi insertions, at cytological locations 30B and 40D, which we verified by genomic PCR. Representative images (A), EdU quantification (A’). n = 6 discs. **(B-B’)** Same as panel (A) except that the second insertion at the 40D locus was recombined away, assessed by genomic PCR. Knockdown with this line also rescues proliferation termination in Tub^ts^>*spok^i^* discs. Representative images (B), EdU quantification (b’). n = 8 discs. **(C-C’)** An independent TSC2 RNAi line from the TRiP collection (Bloomington stock 31770) (Tub^ts^>*spok^i^* + *TSC2*^i^ Trip) also rescues proliferation termination. Representative images (C), EdU quantification (C’). n = 10 discs.

**Figure S9:**
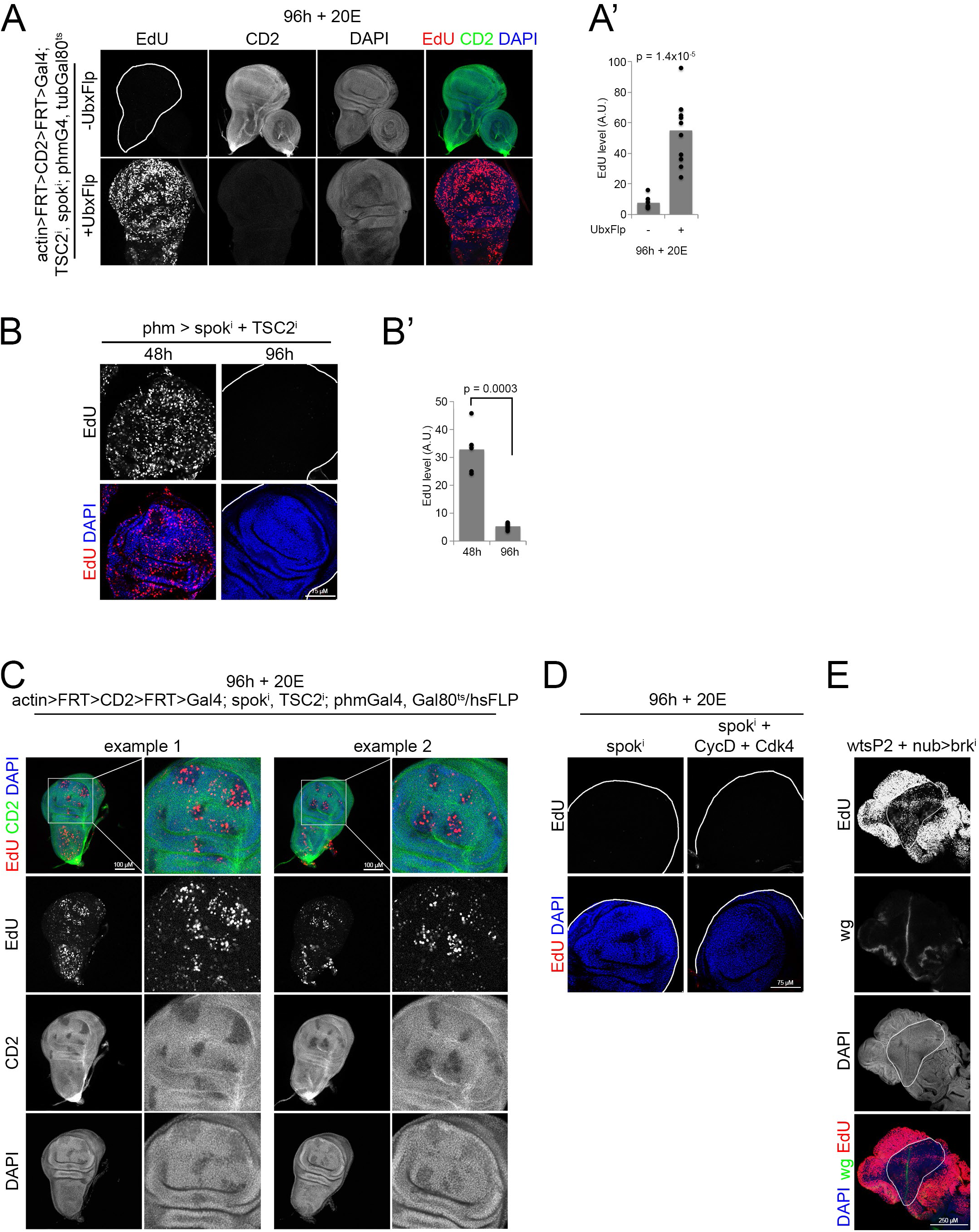
Manipulation of TORC or Cdk4 activity in wing discs at proliferation termination. **(A-B’)** TSC2 knockdown in imaginal discs with UbxFLP bypasses the termination of proliferation of phm^ts^>*spok^i^* +20E discs. (A-A’) Knockdown of TCS2 in both the prothoracic gland and imaginal discs using a combination of phmGAL4 and UbxFlp bypasses the termination of proliferation. Lack of staining anti CD2 shows efficient activation of actin-Gal4 driving TSC2^i^ expression in the presence of UbxFlp. Representative images (A), EdU quantification in (A’). n = 10 discs x 2 replicates. (B-B’) As a negative control, knockdown of TSC2 in the prothoracic gland using phantom-GAL4 (phm^ts^>*spok^i^* + *TSC2*^i^) does not prevent proliferation termination 96h after knockdown induction and 20E feeding. Representative images (B), EdU quantification (B’). n = 6 discs. **(C)** Clones in the wing disc expressing TSC2 RNAi bypass the termination of proliferation at 96h after spok knockdown and 20E feeding. *TSC2^i^* expression was induced by heat shock (45 min, 32°C) and discs were stained for EdU and CD2 (marker for the FLP-mediated recombination event; CD2 negative cells express *TSC2^i^*). n = 36 discs. **(D)** Overexpression of Cyclin D + Cdk4 does not cause a bypass of proliferation termination 96h after spok^i^ induction and 20E feeding. n = 9 discs. **(E)** The pouch (white outline, identified by wg staining) of wts^P2^ discs expressing brk^i^ under the control of nubbin-Gal4 terminate proliferation indicating that brk^i^ is dispensable for proliferation termination in the wing pouch also in the wts^P2^ system. n = 7 discs.

**Figure S10:**
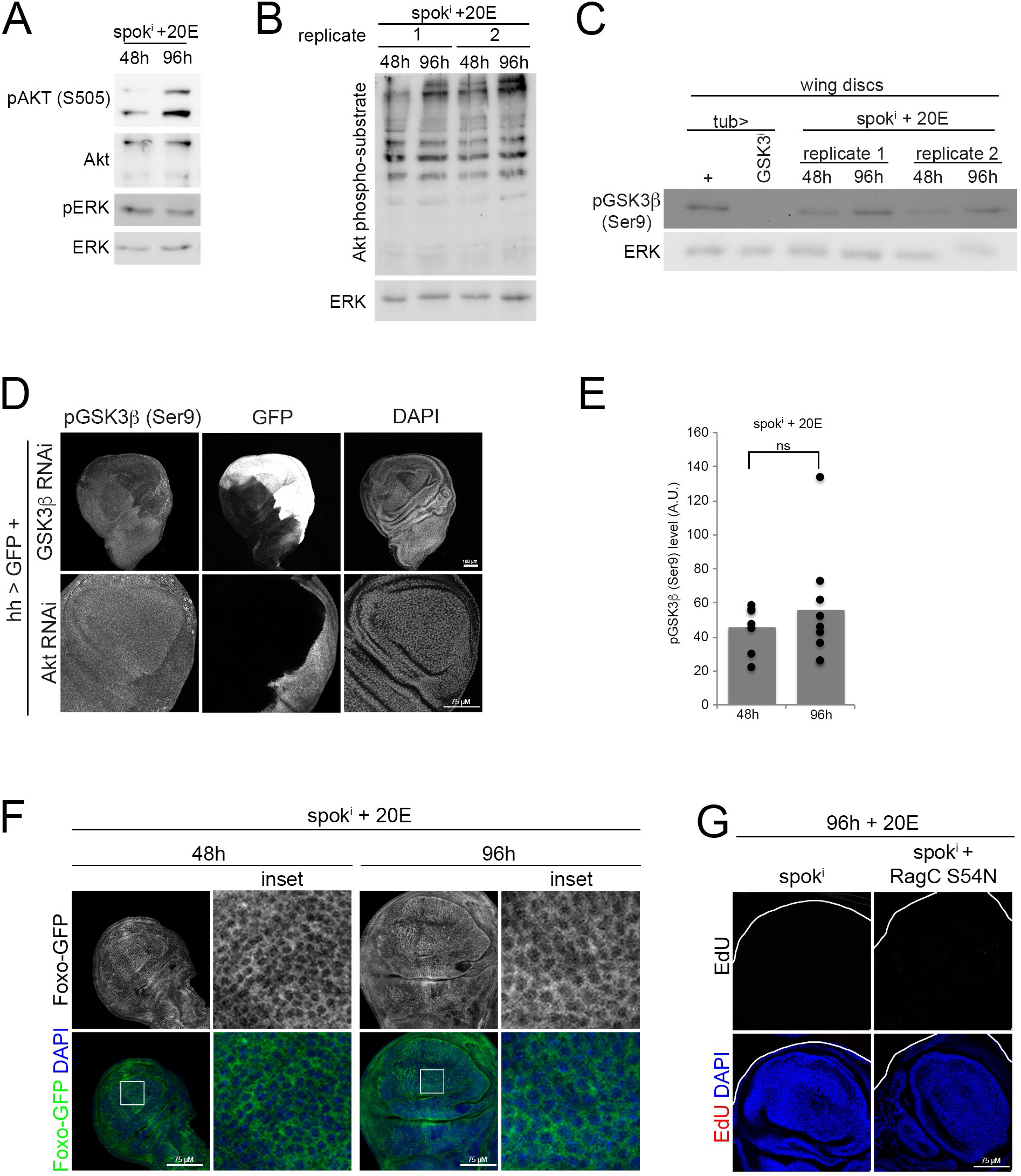
Akt activity does not drop when Tub^ts^>*spok^i^* +20E discs terminate proliferation. **(A-C)** Phosphorylation levels of ERK and Akt (TORC2 site Ser505) (A), of Akt substrates (B) and of GSK3ß (C) were determined by immunoblotting lysates of wing discs from Tub^ts^>*spok^i^* +20E (*spok^i^*) animals either 48 or 96 hours after spok^i^ induction. In (C), knockdown of GSK3ß with Tub-G4 was used as a control to verify specificity of the detected band. n=3 biological replicates x 20 discs/sample. **(D)** The phospho-GSK3ß antibody specifically detects Akt dependent GSK3ß phosphorylation by immunostaining in wing discs, since pGSK3ß staining levels are reduced when either GSK3ß (top panel) or Akt were knocked-down in the posterior compartment using hedgehog(hh)-Gal4. GFP indicates hh expression domain. n = 5 discs. **(E)** Phosphorylation levels of GSK3ß were determined by immunostaining wing discs from Tub^ts^>*spok^i^* +20E animals either 48 or 96 hours after knockdown induction and quantifying integrated density normalized to wing disc area. n = 8 discs. (F) Foxo localization does not change when Tub^ts^>*spok^i^* +20E discs terminate proliferation. Foxo-GFP remains cytoplasmic at 48 and 96 hours after knockdown induction. n = 4 discs. **(G)** Overexpression of constitutively active RagC does not bypass proliferation termination. Wing discs from Tub^ts^>*spok^i^* +RagC^S54N^ +20E larvae stop proliferating 96h after knockdown induction and 20E feeding. n = 13 discs.

**Supplemental Table 1.**
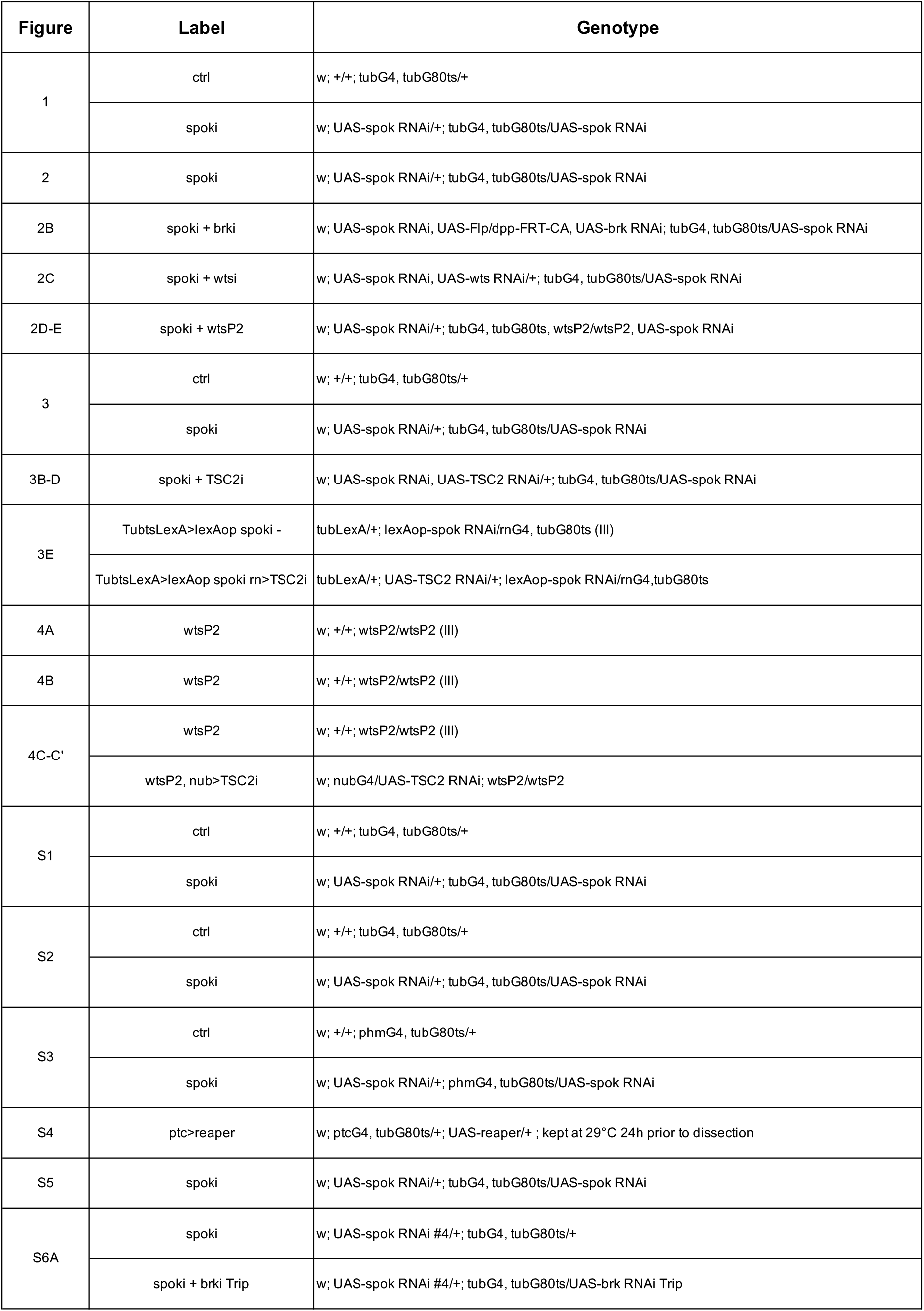
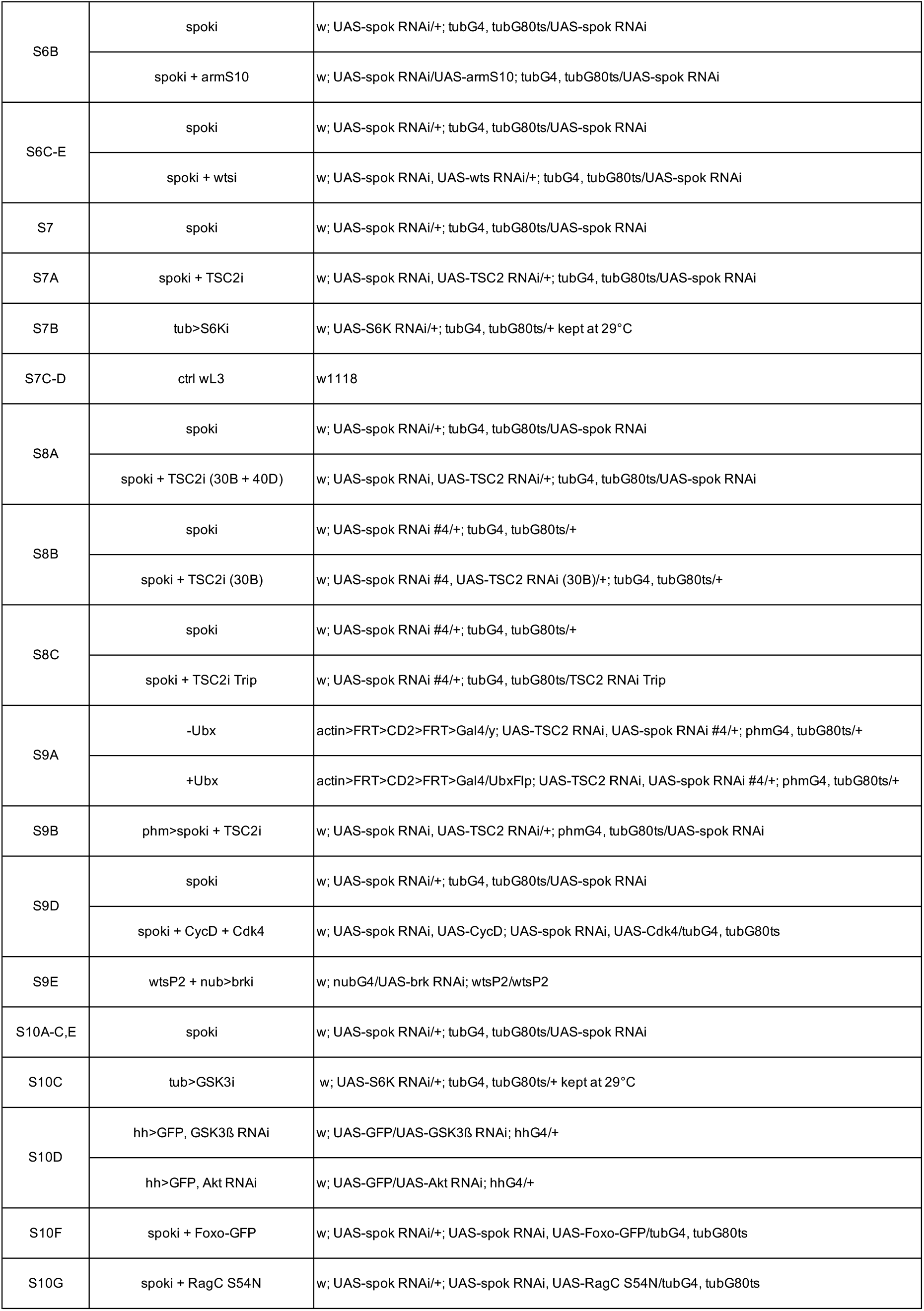
genotypes.

## Notes

### Competing Interest Statement

The authors have declared no competing interest.

